# Unraveling the complexity of rat object vision requires a full convolutional network - and beyond

**DOI:** 10.1101/2024.05.08.593112

**Authors:** Paolo Muratore, Alireza Alemi, Davide Zoccolan

## Abstract

Despite their prominence as model systems to dissect visual cortical circuitry, it remains unclear whether rodents are capable of truly advanced processing of visual information. Here, we considered several psychophysical studies of rat object vision, and we used a deep convolutional neural network (CNN) to measure the computational complexity required to account for the patterns of rat performances reported in these studies, as well as for the animals’ perceptual strategies. We found that at least half of the CNN depth was required to match the modulation of rat classification accuracy in tasks where objects underwent variations of size, position and orientation. However, the full network was needed to equal the tolerance of rat perception to more severe image manipulations, such as partial occlusion and reduction of objects to their outlines. Finally, rats displayed a perceptual strategy that was way more invariant than that of the CNN, as they more consistently relied on the same set of diagnostic features across object transformations. Overall, these results reveal an unexpected level of sophistication of rat object vision, while reinforcing the intuition that, despite their proficiency in solving challenging image classification tasks, CNNs learn solutions that only marginally match those of biological visual systems.

## Introduction

The neuronal basis of visual object recognition has been the subject of intense study, with a particular focus on the ventral visual pathway - the visual cortical hierarchy that, in primates, is devoted to process object and shape information^1,2^. Recent behavioral and neurophysiological evidence suggests that also rodents, and rats in particular, display nontrivial object recognition abilities^3^, thus offering a viable new model for studying the neuronal basis of object vision. Rats are able to recognize objects despite variations in pose, size, position, illumination and visual cues^4–7^. They can discriminate the content of natural movies^8^ and even categorize faces^9^. In addition, rats display the same pattern of sensitivity to multi-point texture statistics that humans do^10^ and they seem to process visual objects by relaying on nontrivial, multifeatural perceptual strategies^5,9,11,12^. These perceptual abilities are consistent with the hierarchical increase in the complexity and invariance of object representations found along the progression of extrastriate visual areas that run laterally to primary visual cortex (V1), pointing to this pathway as the rat homologue of the primate ventral stream^7,13–17^.

Despite the behavioral and neurophysiological evidence reviewed above, a limit of current visual perceptual studies in rodents is the relatively low number of visual stimuli they rely upon, which rarely exceeds a hundred of distinct object conditions (but see refs.^5,9,11,12^). By comparison, modern investigations of primate object vision benefit from high-throughput psychophysical techniques that allow testing at a much larger scale (i.e., order of thousands of images across hundreds of human participants)^18,19^. The limited size of stimulus sets used in rodent studies makes it hard to fully exclude the possibility that the animals succeed in a given discrimination tasks by relying on trivial strategies based on detection of low-level visual cues. For instance, we have previously shown that, when objects are presented across a limited number of transformations (or views), differences of mean luminosity between the sets of views of two objects allow invariant encoding of object identity already at the level of V1 representations^7^.

One way to check whether the variety of stimulus conditions employed in a object recognition task was large enough to engage high-order representations is to feed the same battery of conditions to a Convolutional Neural Network (CNN) for image classification and then measure how well the task is solved across consecutive layers of the network. This approach relies on the fact that CNNs, with their hierarchical architecture and ability to learn complex representations, have proven to be the most successful artificial models for biological vision to date^20–24^. In particular, several studies have found a hierarchical match between ventral stream areas and CNN layers in terms of the ability of neural activations in the latter to explain neuronal tuning in the former. That is, while activations in early CNN layers better predict responses to natural images in low-level visual cortical areas (e.g., V1), activations in deeper layers better account for neuronal tuning in higher stages of the ventral stream, such as the inferotemporal cortex (IT)^21,25,26^. This suggests that a similar approach could be used to assess the complexity of an object recognition task, by measuring at which depth of a CNN object representations successfully solve the task. Trivial discriminations would be solvable by represenations in early layers, while more demanding tasks (e.g., in terms of invariance) would require the full extent of the neural architecture.

This procedure has been applied to assess the complexity of visual representations used by humans engaged in a rapid animal vs. non-animal categorization task^27^. More recently, a first attempt to gauge the complexity of rat object vision using this approach was presented by Vinken and colleagues^28^. The authors found that rat classification accuracy reported in several object recognition experiments^4^ was successfully modelled by very early layers of standard CNNs - more computational depth was required only when the task involved discrimination of natural movies^8^. In a follow-up study^29^, CNNs were used to devise a suitable image set to investigate the extent to which rats are capable of truly advanced object recognition. Although rat classification accuracy was best captured by activations in late convolutional layers of a CNN, the latter could only account for a small fraction of the variance of rat behavioral performance and substantial differences emerged with the performance pattern of human participants tested in the same task.

Overall, these results suggest that former psychophysical studies may have overestimated the ability of rats to tolerate transformations in object appearance, because the size and variety of the stimulus sets may not have been sufficiently large to rule out the deployment of low-level strategies. However, we believe that previous applications of CNNs to evaluate the complexity of rat vision lacked important details concerning the resolution and noise at the front end of the rat visual system, the way in which the animals viewed the stimuli in the absence of head fixation, and the cognitive constraints that affect perceptual decision making in rodents. To overcome these shortcomings, we developed an image pre-processing pipeline that explicitly models the constraints of rat vision (e.g., low visual acuity^30,31^) and the additional image variability (e.g., translations and in-plane rotations) induced by rat head movements during execution of the perceptual task^32^. Furthermore, we went beyond a comparison between rats and CNNs in terms of absolute discrimination performances - which are not that meaningful, considering that the large lapse rates of rodent perceptual decisions^33^ are fully absent in CNN models. We focused instead on comparing how similarly recognition accuracy was modulated across object views in rats and CNN layers. In addition, we compared the visual strategies used by rats and CNNs to solve the same object recognition tasks, by applying to CNNs the image classification approaches that have been successfully used to infer rat visual perceptual templates in refs.^5,12^. This approach aligns with recent strides in the field of explainable artificial intelligence^34^ and cognitive sciences^35^ that have highlighted how modern architectures for machine vision, despite their saturating classification accuracies on challenging benchmarks, often exploit unintelligible visual strategies^36^ (e.g., features in the background) that substantially differ from those used by their biological counterpart^37,38^.

We found that, although mid-level layers of a standard CNN (VGG-16) are effective models of rat classification accuracy in tasks with moderate image variability, the full depth of the network, including the final, fully connected layers, is required to explain rat performance on more challenging tasks, such as those involving partial occlusions or outline versions of the target objects. In addition, we found that rat visual perceptual strategies are more invariant than those used by the CNN. Rats make more consistent use of the same set of visual features, preserved across transformations, thus relying on a more parsimonious and generalizable collection of visual landmarks than the corresponding artificial counterpart. Interestingly, these findings are consistent with several of the discrepancies observed between humans and CNNs in terms of image processing (see Discussion). This reasserts the sophistication of rat vision and the potential of rodents as model systems to investigate nontrivial visual computations. At the same time, it suggests possible adjustments to the architecture and visual diet of CNNs to improve their object awareness^39^ and their robustness to out-of-distribution image manipulations in which both primate and rodent visual systems appear to excel.

## Results

Our goal was to establish the extent to which rats are capable of advanced shape processing by comparing their performance patterns and perceptual strategies, as reported in three of our previous studies^4,5,12^, with those afforded by object representations across the layers of VGG-16 - a popular convolutional network that has been often used as a benchmark against which to compare biological vision (e.g., see refs.^26,28,40–44^). This was achieved by feeding the CNN with the same object conditions employed in the three rat studies, using the data processing pathway that is depicted in Figure 1.

**Figure 1:**
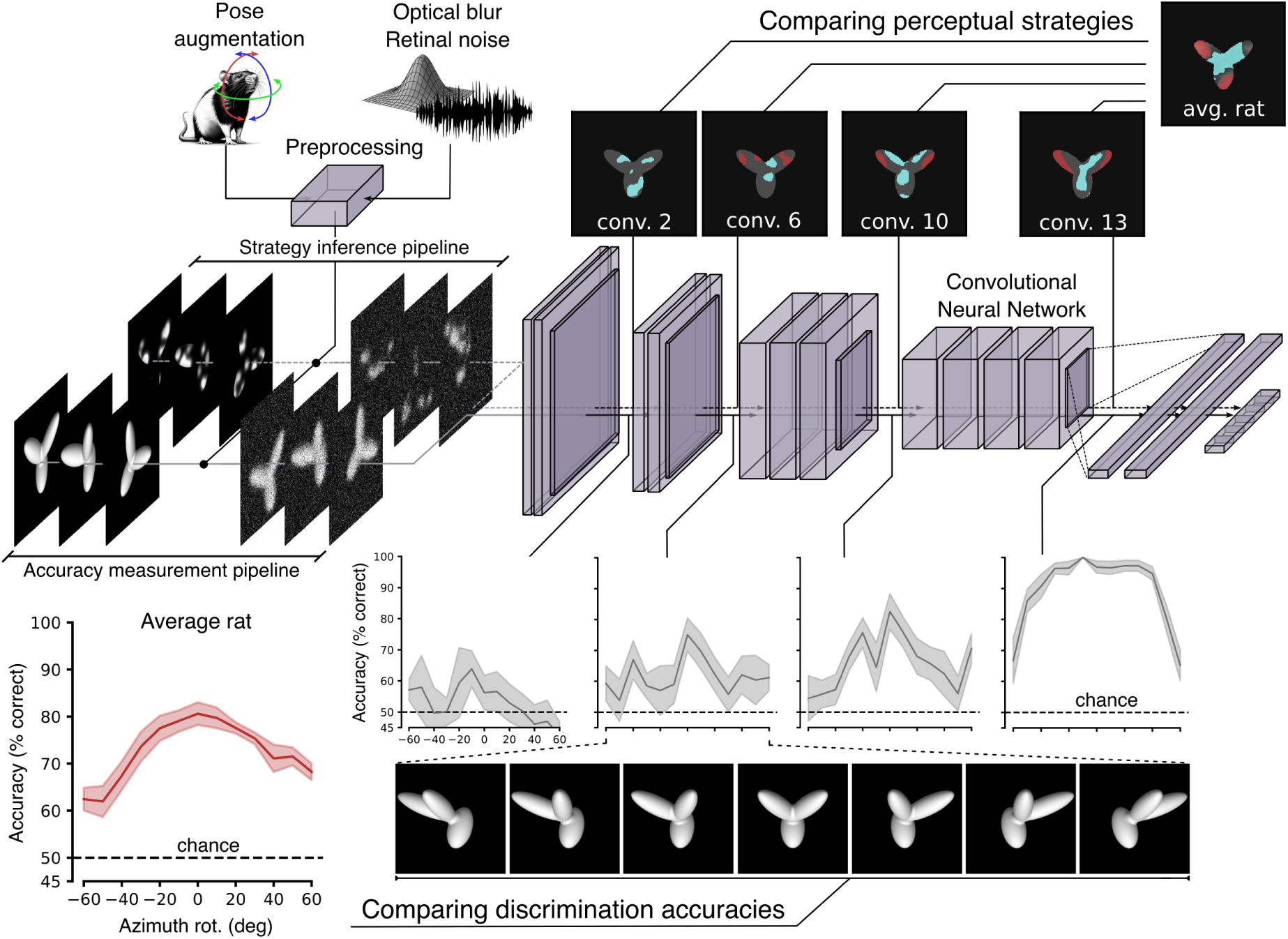
Comparing processing of visual objects in rats and CNNs. Conceptual overview of the experiments we carried out on VGG-16 to assess the complexity of rat object vision. We investigated how consecutive layers of VGG-16 represent the visual stimuli that have been used in three previous studies of rat visual object recognition^4,5,12^. Before being fed to VGG-16 input layer, the visual objects were subjected to an image pre-processing stage to simulate the resolution/fidelity of rat spatial vision (by the addition of blur and noise), as well as the variations in object appearance produced by head movements (resulting in an augmented image set). Following the experimental design of the rat studies, visual objects were presented to the network in two different flavors. Unoccluded/unaltered images were used to measure how well object representations across VGG layers supported object discrimination (bottom branch: *accuracy measurement* pipeline). This allowed comparing how well the pattern of rat discrimination accuracies across object transformations (e.g., azimuth rotations; red curve) aligned to the patterns of discrimination accuracies measured along the network (gray curves). Conversely, either partially occluded or structurally altered versions of the target objects were used to infer what visual strategies VGG representations afforded to support object discrimination (top branch: *strategy inference* pipeline). This allowed comparing the set of diagnostic salient (red patches) and anti-salient (cyan patches) features used by the rats (rightmost saliency map) and by VGG (sequence of saliency maps along checkpoint layers) to succeed in the object recognition task. For both pipelines, VGG layers were probed by training linear classifiers to predict the image labels based on the object representations they provided.

Since our aim was to measure both the object discrimination accuracy and the underlying perceptual strategy, and since the latter was uncovered, in the rat studies, using two different classification image approaches (where the original objects were altered either by partial occlusion with random masks or by random structural variations), the visual objects fed to the network were either the unoccluded/unaltered objects (*accuracy measurement* pipeline in Fig. 1) or their occluded/structurally altered versions (*strategy inference* pipeline). In both cases, before being fed to VGG input layer, the images of the visual objects were subjected to a pre-processing step to augment the original stimulus dataset in such a way to: 1) better match it to the low resolution of rat spatial vision (i.e., by adding blur and noise); and 2) simulate the additional variations in object appearance induced by head movements^32^, given that the animals were not head-fixed but only partially body restrained.

We then followed an approach that is similar to the one adopted by previous studies comparing CNNs’ with humans’ or rats’ proficiency in object recognition tasks^27,28^. We used linear classifiers to read out the identity of the objects fed to VGG-16, based on their representations in each layer of the network. This method allows probing the extent to which object representations at a given stage of a processing hierarchy are sufficiently untangled to support transformation-tolerant recognition by a simple, linear readout scheme^2,45^. By systematically and independently applying a linear decoder to each layer of VGG-16, we could thus scan the depth of the network, pursuing two complementary goals: 1) to look for the processing stages yielding the patterns of discrimination accuracy that were most consistent with those measured for the rats (*accuracy measurement* pipeline: see the bottom branch of Fig. 1); or (2) to infer the visual discrimination strategies supported at each stage of the network and compare them to the known perceptual templates deployed by the rats (*strategy inference* pipeline: see the top branch of Fig. 1).

More specifically, we trained a linear Support Vector Machine (SVM) per layer on the object classification task, using, as feature vectors, the activations of a random sub-population of units for the training views of the two objects. In any given experiment, the size of the sub-populations sampled from each convolutional layer was held constant, so as to control for the possible impact of the dimensionality of the representational space on classification accuracy. In addition, since the layers widely differed in the total number of units, we further controlled for potential population under-sampling by repeating each test with an increasing (logarithmically spaced) number of units (from 10^3^ to 10^5^), capping them to the total size of a layer when needed (i.e., in the fully connected layers). Finally, each trained SVM was tested for its ability to correctly classify the activations produced by the held-out test images and both the train and test accuracies were recorded.

### Image augmentation pipeline to account for rat visual acuity and head movements

A fair comparison between rat visual perception and CNN models requires the latter to incorporate some key ecological constraints on the native resolution of the rat visual system. Rats have low visual acuity, being able to resolve at most a spatial frequency (SF) of 1 cycle/deg^30,31,46^. This imposes a filter on the quality of the visual information that the front-end of the rat visual system conveys to the rest of the brain. Spatial acuity is typically quantified via the Contrast Sensitivity Function (CSF)^30^. The CSF is inversely related to the minimal amount of contrast that is required for successful detection of a sinusoidal grating at a given spatial frequency (see Methods). In the case of Long-Evans rats, the curve peaks at 0.1 cycles/deg and drops to zero for SFs smaller than 0.04 cycles/deg and higher than 1 cycles/deg (black curve in Fig 2a). This shape results from the interplay of several factors, such as the quality of the optics of the eye, the density of cones and rods in the retina, and the level of neuronal noise in the photoreceptors and the other retinal cell types^47–49^. All these factors contribute to determine the quality of the retinal image, which needs to be taken into account in the assessment of rat pattern vision. Since developing an anatomical and biophysical model of the encoding of retinal images would be a daunting task, also because of the paucity of available data, we resorted to a functional modelling approach. Considering the CSF as the aggregate functional measurement of the sharpness/resolution and fidelity of rat spatial vision, we searched for the combination of image blurring and noise that best reproduced the rat CSF.

**Figure 2:**
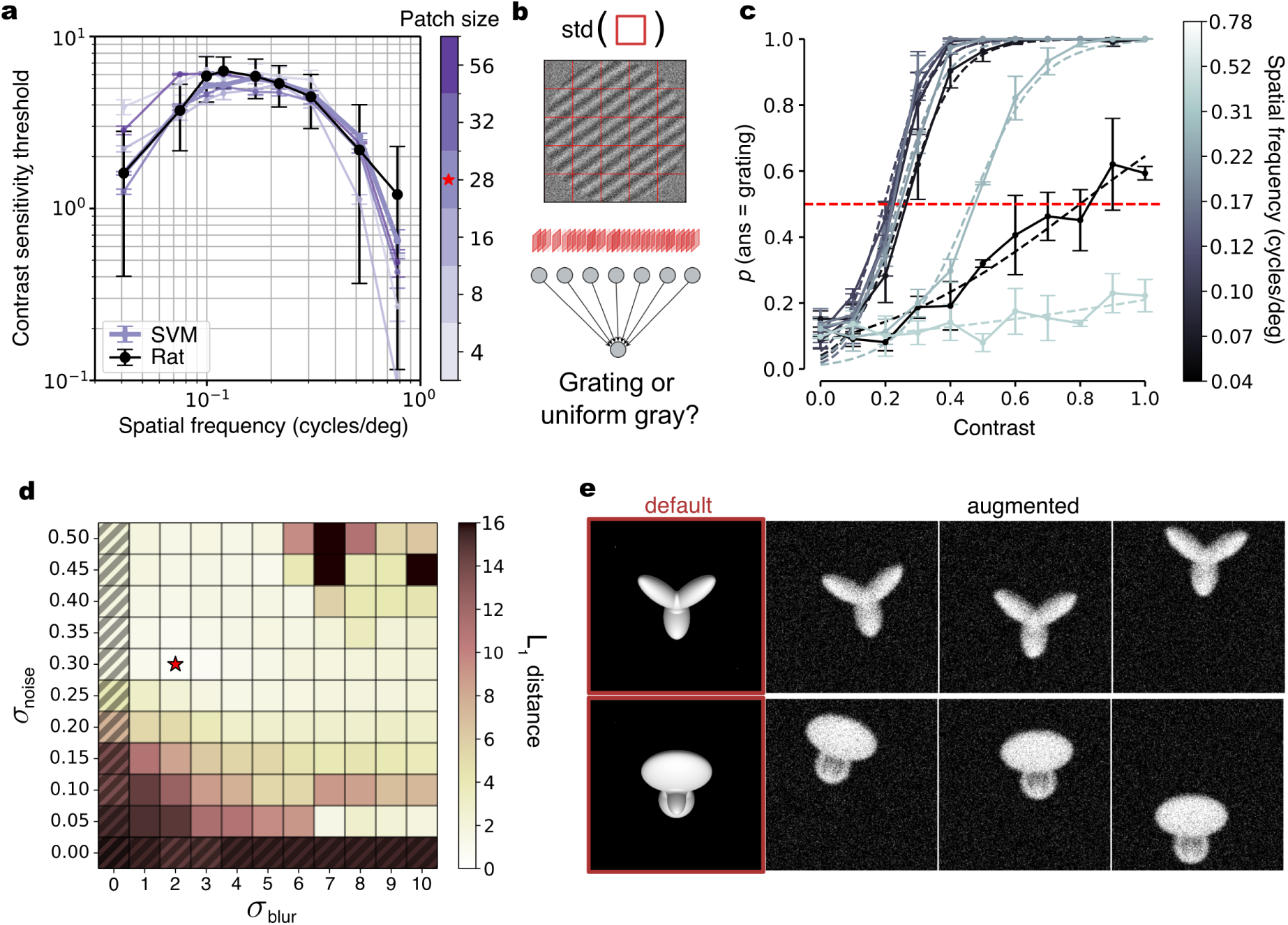
Image augmentation pipeline to account for rat visual acuity and head movements. (**a**) The Contrast Sensitivity Function (CSF) measured for rats by Keller^30^ (black dots/line) is shown along the CSFs (purple curves) obtained for a simulated observer, measuring the contrast of input images in small patches with different sizes (as illustrated in **b**). The curves all refer to the same combination of image blur and noise marked by the red star in **d**. The red star on the color bar marks the patch size yielding the CSF with the best fit to the rat CSF. (**b**) Schematic representation of the grating detection task used to obtain the CSFs of the simulated observer. The task consisted in detecting whether the input image was a grating, as opposed to a uniform gray square, by relying on the contrast measured over a grid of patches within the image (red squares). The images were subjected to various degrees of Gaussian blur and additive noise. The simulated observer consisted in a linear classifier, fed with the contrast values measured across the image patches. (**c**) Psychometric-like functions, showing the probability of the simulated observer to detect an input grating (with a given spatial frequency) as a function of its contrast. The contrast level at which a given psychometric crossed the *p* = 0.5 value was taken as the contrast threshold for that particular spatial frequency. Such thresholds were used to obtain the CSFs, as explained in the Methods. The curves shown here were obtained for the combination of parameters marked by the red star in **d** and path-size = 32. (**d**) *L*_1_ distance between the rat CSF and the CSFs of the simulated observer that were obtained over a combination of *σ*_noise_ and *σ*_blur_ parameters for patch-size = 28. The red star marks the combination yielding the best fit (same for all tested patch sizes). (**e**) Outcome of the full augmentation pipeline applied to the default views of the objects used in refs.^4,5^. For each of the original objects (highlighted in red), three augmented views are provided, resulting from adding the blur and noise levels yielding the best fit with rat CSF and from applying the expected vertical/horizontal shifts and in-plane rotations produced by head movements.

To simulate the grating detection task used to measure the CSF, we trained a linear Support Vector Machine (SVM) to discriminate noisy and blurred sinusoidal gratings from noisy, uniform middle gray images. More specifically, we divided the simulated stimulus display in a grid of squared patches, we measured the image contrast within a patch as the standard deviation of its pixel intensity values and we took the resulting set of patch contrasts as the feature vector to be fed to the SVM (Fig. 2b). Multiple samples (5,000) for each class of stimuli (gratings and middle gray images) were produced by random variations of the additive noise pattern applied to the images and, in the case of the gratings, by random variations in phase and orientation.

This pipeline simulated an observer that tries to detect the presence of a grating based on the luminance contrast extracted from the stimulus display. This allowed measuring the detection performance of the simulated observer as a function of image contrast and spatial frequency, thus obtaining psychometric functions from which we could estimate the contrast thresholds to detect the gratings at each SF (Fig. 2c). Based on these thresholds, we could compute the CSF for the simulated observer and compare it to the rat experimental CSF measured by Keller^30^ and reported in Figure 2a (back curve). To estimate the blur and noise levels that yielded the closest match (i.e., lowest *L*_1_ distance) between simulated and experimental CSFs, we performed a grid-search over the Gaussian blur intensity and the Gaussian noise applied to the input images (quantified via their variance 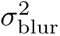 and 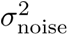, respectively), as well as over the patch size *p* (Fig. 2d). The resulting *L*_1_ landscape over the 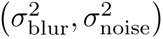 plane featured a flat valley (light cells) with the absolute minimum (marked by the star) yielding a very precise fit with the experimental CFS regardless of patch size (Fig. 2a; compare the purple curves to the black curve). Thus, with a moderate level of blur and noise, we could functionally and accurately simulate the quality of the images encoded by the rat retina and use this preprocessing step in all our tests with VGG-16.

A second preprocessing stage was also applied to take into account the fact that all perceptual studies of rat object vision to date have been carried out without head fixation and gaze control^3^. In the experiments performed by our group, some level of body restraint has been achieved by requiring the animals to protrude the head through a narrow viewing hole placed in front of the stimulus display^4,5,10–12,50,51^. This allows for a good control over viewing distance and, therefore, stimulus size, but does not fully prevent head movements. In particular, we found that head orientation at the time of stimulus presentation is not reproducible across trials: pitch, roll and yaw rotations vary over a span of about 60°, 35° and 20°, respectively^32^. Obviously, these trial-by-trial variations in head orientation induce additional transformations on the images of the visual stimuli over the rat retina, on top of the image variability designed by the experimenters. Specifically, pitch and yaw rotations translate into vertical and horizontal shifts, while roll rotations produce in-plane rotations. With knowledge of the geometry of the experimental rig and basic trigonometry, one can easily compute the corresponding image transformations (see Methods). These can be considered as augmented versions of the training and test images used to probe object recognition in rats. As an example, in Figure 2e we have reported the effects of the described augmentation on the images of the original stimuli used in refs.^4,5^. The default views of the objects used in those studies are shown on the left (highlighted by the red frames), while a few instances of augmented versions (incorporating also the blur and noise produced by the preprocessing step described above) are displayed on the right.

Since, in a given trial, head orientation along the three rotation axes was random (within the ranges reported above), to properly simulate the actual level of image variation experienced by the rats during the training/testing in our perceptual tasks, we implemented the following pipeline. For each visual image that had to be fed to the CNN, we first randomly sampled the values of pitch, roll and yaw within the allowed ranges, then we translated them into the corresponding vertical/horizontal shifts and in-plane rotations of the image. After this augmentation step was completed, we added the blur and random noise perturbations and we finally fed the image to the network. For brevity, in what follows, we will refer to this whole preprocessing pipeline (including augmentation, blurring and noise) simply as “augmentation”. The pipeline was consistently applied to all the images fed to the CNN that are described in our study, as illustrated in Figure 1.

### Half of VGG-16 depth is needed to match rat tolerance to size changes and in-depth rotations

As a first assessment of the complexity of rat object vision, we considered the experiment originally presented by Zoccolan and colleagues^4^, who tested the ability of rats to discriminate two visual objects despite variations in size and in-depth rotations (the two visual objects are shown in Fig. 3a, left; the full set of transformations applied to one of the objects is shown in Figure 3a, right). The rats were initially trained to discriminate two specific (”frontal”) views of the objects (those shown in Fig. 3a, left), presented at 40° of visual angle (blue frame in Fig. 3a, right). Then, they were trained on scaled versions of the frontal views (down to 15° of visual angle) and on in-depth, azimuth rotations of the objects (from −60 to + 60 degrees) shown at 30° of visual angle (these further training conditions are indicated by the red cross in Fig. 3a, right). Finally, the rats were tested on unseen combinations of size and azimuth transformations (off-cross conditions in Fig. 3a, right). The animals achieved above-chance discrimination accuracy across virtually all tested conditions, although the performance decreased as a function of the magnitude of the transformation (Fig. 3b).

**Figure 3:**
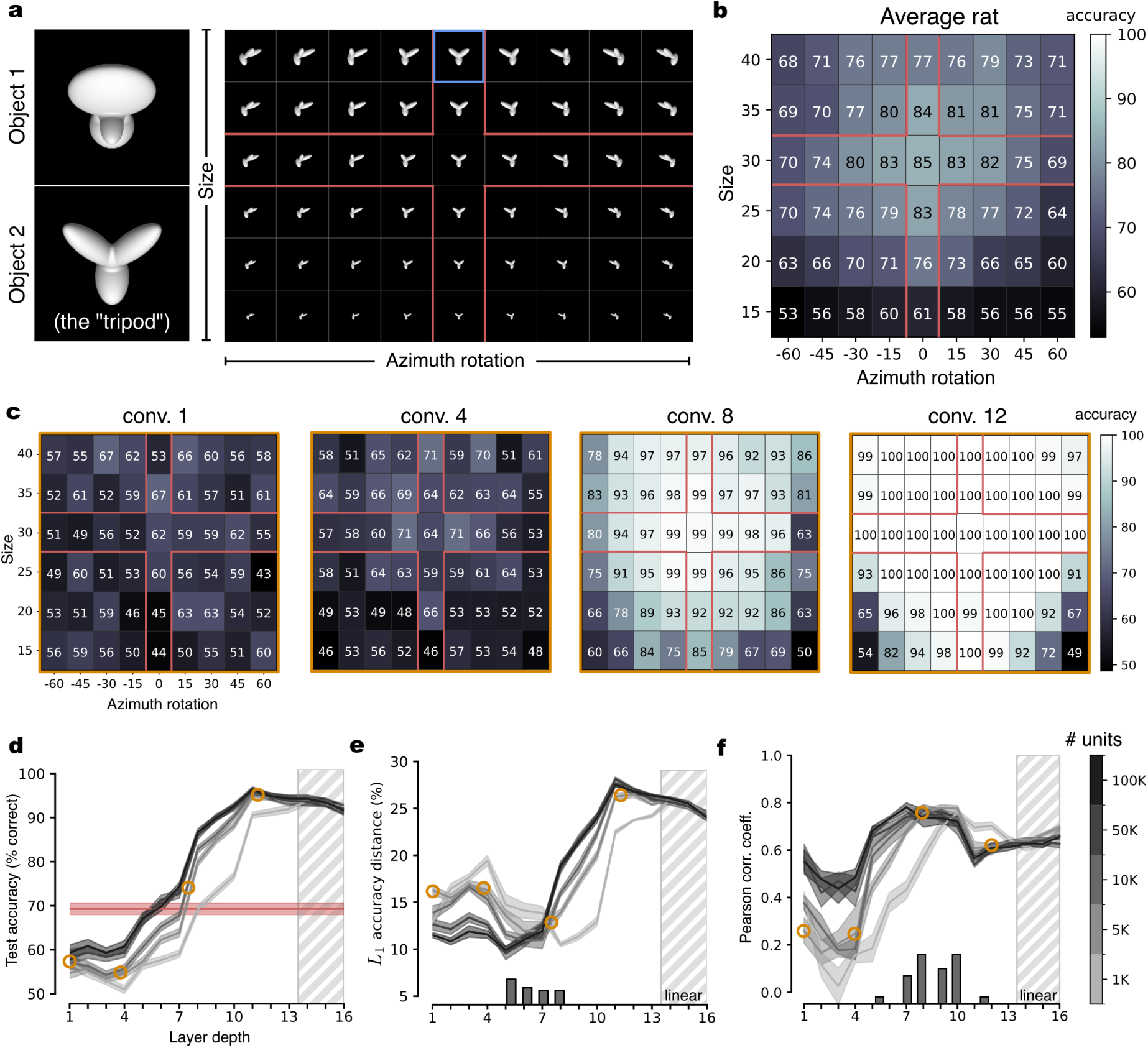
VGG-16 mid-level layers match the pattern of rat discrimination accuracy across size changes and in-depth rotations. (**a**) The stimulus set originally used by Zoccolan and colleagues^4^. Two objects (default views shown on the left) were presented across a combination of size and azimuth in-depth rotations (shown on the right for one object). The blue frame indicates the view used for the initial training of the animals. The red cross indicate the additional views the rats were trained with, before being tested with the off-cross views. (**b**) Group average performance of the rats across the set of object transformations shown in **a**. Adapted from ref.^4^. (**c**) Discrimination performances achieved by training SVM classifiers with the activations of VGG-16 units to the on-cross object views and testing them with the activations to the off-cross views of the stimulus matrix shown in **a**. The four performance matrices refer to activations sampled from progressively deeper convolutional layers of the network. (**d**) Test accuracies (i.e., performances measured on the off-cross views of **a**) of the SVM classifiers achieved across the depth of VGG-16 (gray curves). Shaded areas are SEM over five different classification runs (see Methods). The shades of gray indicate the number of units that were sampled from each layer and fed to the SVMs (see colorbar on the right of **f**). Rat average test accuracy is reported in red. The orange circles highlight those layers, whose accuracy patterns have been displayed in **c**. (**e**) *L*_1_ distance between the pattern of accuracy of the rats across the whole stimulus matrix (i.e., both on- and off-cross cells in **b**) and the patterns of accuracy obtained across VGG-16 layers. The histogram in the bottom reports the occurrence of the minima of the *L*_1_ distance across classification runs and sizes of the populations fed to the SVMs. Cyan circles and shaded areas as in **d**. (**f**) Same analysis as in **e**, but carried out by computing the Pearson correlation coefficient instead of the *L*_1_ distance.

To check the extent to which this performance pattern is consistent with advanced processing of object information, we administered the same task to VGG-16, following the *accuracy measurement* pipeline illustrated in Figure 1. We used VGG-16 pretrained to achieve high classification performances on Imagenet, which ensured that populations of units across progressively deeper stages of the network represented increasingly complex combinations of features found in natural images in an increasingly transformation-tolerant way^52–54^. We fed the network with thousands of randomly augmented versions (see previous section) of the two objects used in the rat experiment, sampled from the whole pool of transformations (i.e., both on- and off-cross views in Fig. 3a, right). Following the approach described in Figure 1, we trained linear SVMs on the object discrimination task using train examples only (i.e., on-cross images in Fig. 3a, right) and we tested the classifiers on held-out views (i.e., off-cross images), looking for the processing stages that yielded the pattern of discrimination accuracy that was most consistent with the one measured for the rats (shown in Fig. 3b).

Figure 3c reports the patterns of discrimination accuracies of the SVM classifiers over the combination of size and azimuth transformations that we measured across progressively deeper checkpoint layers of VGG-16. A qualitative inspection of these patterns shows a progression from near-chance performances in initial layers towards saturating, near-perfect performances in the final ones. This trend was quantified in Fig. 3d, which reports the average performance (gray lines) across all object views included in the test set (i.e., off-cross cells in Fig. 3a, right), along with the average performance measured for the rat on those same conditions (red line). Discrimination accuracy started close to chance and remained such until layer 4, where it started to increase sharply, crossing rat performance between layer 6 and 8, depending on the size of the sub-populations of units fed to the SVMs. Larger sub-populations (darker curves) yielded higher performances, although the difference was mainly appreciable when the number of units increased from 1K to 5K, while further expansions till 100K only added a marginal gain. Performance saturated near to 90% correct in the deepest layers.

A finer-grain comparison between the magnitude of rat and VGG performances was obtained by computing the *L*_1_ distance between the performance matrix of the rats (i.e., the data shown in Fig. 3b) and the performance matrices obtained across consecutive layers of the network (like the examples shown in Fig. 3c). The smallest distance was observed between layers 5 and 8 (Fig. 3e), as shown by the distribution of the *L*_1_ minima obtained across 5 different classification runs for each sub-population size (histogram at the bottom of Fig. 3e).

Overall, according to these analyses, it takes between 6-8 convolutional layers of a powerful CNN architecture to match rat accuracy on the object recognition task used ref.^4^. This shows that dealing with the level of image variability imposed by the task is not trivial and requires substantial processing. This conclusion is, by itself, in contrast with the one of Vinken and colleaues^28^, who found that VGG-16 first layer already surpassed rat performance on the task (see Discussion). More importantly, as already mentioned in the Introduction, we believe that comparing the rat visual system to CNNs in terms of absolute performances is not very meaningful in the first place. The two systems are too different, not only at the level of the basic architecture (i.e., number of processing stages, number of units, etc) but even in terms of perceptual decision strategies. For instance, rats display large lapse rates, i.e., constant rate of errors even on “easy” stimuli (where performance should be perfect), which do not reflect an inability to correctly recognize the stimuli but, rather, the deployment of exploratory strategies^33^. These lapses impose a cap on the maximal accuracy a rat can reach on a given task, thus leading to a systematic underestimation of the actual perceptual discriminability of the stimuli.

In the light of these considerations, we also carried out a different sort of comparison, measuring the Pearson correlation coefficient between the patterns of discrimination accuracies obtained for the rats and each VGG layer across the transformation matrix. Being scale and shift invariant, Pearson correlation allows assessing the extent to which rat and VGG performances are similarly modulated by the combinations of size and azimuth variations, regardless of the magnitude of the performance. As shown in Figure 3f, Pearson correlations steadily raised across the depth of VGG, reaching a flat peak between layers 7 and 10 (see the histogram of maximal correlations across classification runs and sub-population sizes and at the bottom of Fig. 3f). The magnitude of the correlations at the peaks was substantial (about 0.8), thus confirming the visual impression of a very tight match between the classification performance landscapes obtained for the rats and for middle layers of VGG-16 (compare Fig. 3b to the third matrix in Fig. 3c). This result indicates that, when absolute performance magnitudes are left aside, it takes at least half the computational depth of VGG-16 to closely match rat sensitivity to the variations in object appearance tested by Zoccolan and colleagues^4^.

### The whole depth of VGG-16 is required to match rat robustness to partial occlusion

A limitation of the image set used by Zoccolan and colleagues^4^ (Fig. 3a) is that it included only two kinds of object transformations: size changes and in-depth azimuth rotations. Other studies have probed the tolerance of rat object vision with a wider variety of transformations. In particular, Alemi-Neissi and colleagues^5^ probed rat vision with the same objects used in ref.^4^, but subjected also to horizontal translations and in-plane rotations, in addition to size and azimuth variations. Moreover, rats were also exposed to versions of the objects that were partially occluded by opaque masks punctured by randomly located transparent openings, or “bubbles” (examples of unoccluded and occluded conditions for the two objects are illustrated in Fig. 4a). Overall, this offered the opportunity to investigate the similarity between rat and VGG-16 invariant recognition in the case of a wider, more challenging pool of image transformations.

**Figure 4:**
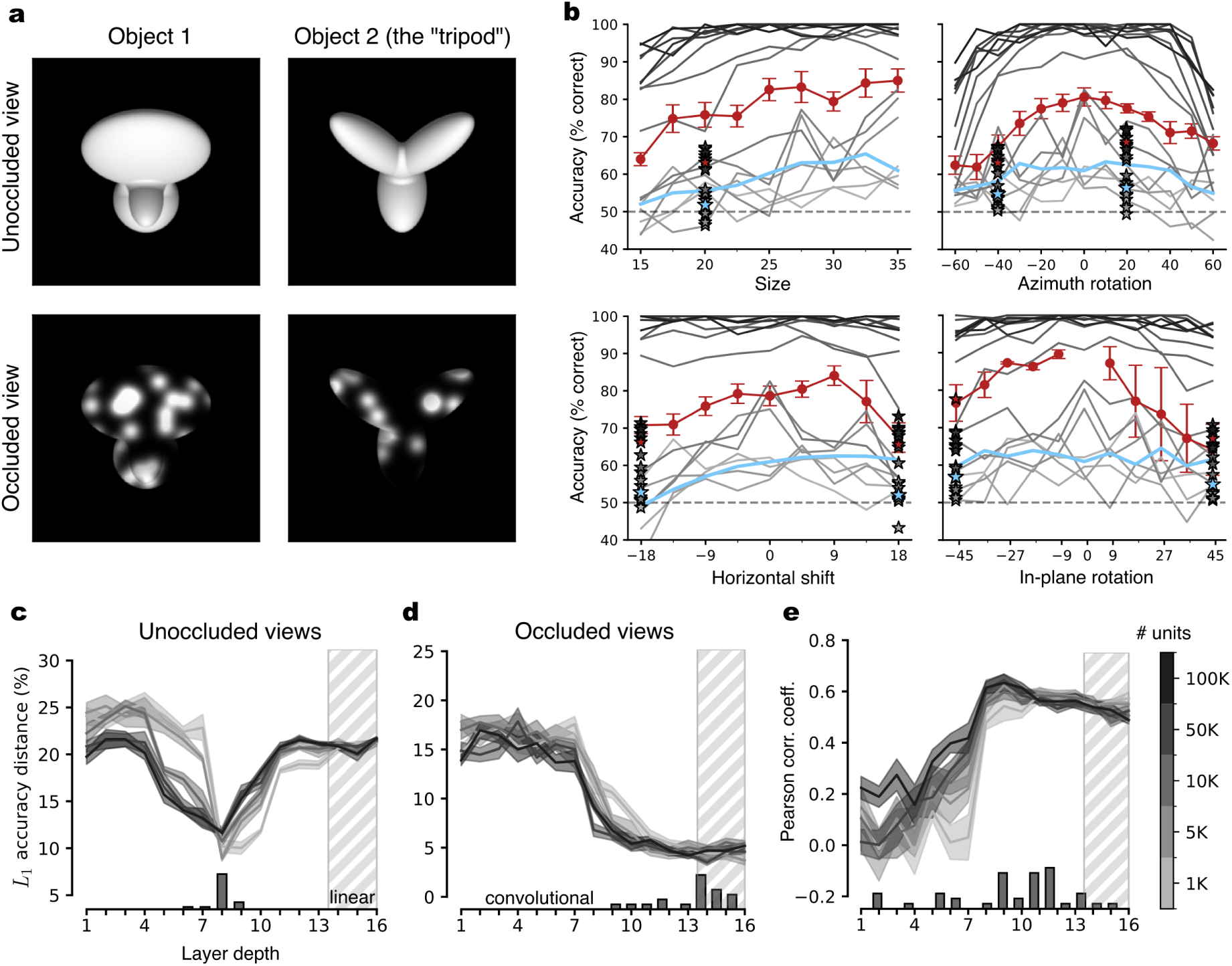
VGG-16 deepest layers are necessary to match rat tolerance to partial occlusion. (**a**) The default views of the two objects used by Alemi-Neissi and colleagues^5^ (top) are shown along two examples of partially occluded views (bottom). (**b**) Classification accuracy of linear SVMs trained to discriminate the two visual objects shown in **a** based on the representations provided by the activations of VGG-16 units across progressively deeper layers (100K units were randomly sampled from each layer). Discrimination accuracy was measured along four different axes of image variation: size, azimuth rotation, horizontal shift and in-plane rotation. The performances of linear SVMs directly trained on the pixel-level representations are shown in cyan. The gray curves report the accuracy measured on unoccluded images, across 5 repetitions of the classification procedure (see Methods). The gray stars indicate the accuracy achieved by the SVMs on a few selected object views that were partially occluded (as shown in **a**, bottom). The shades of gray (from light to dark) code for layer depth (from early to deep). The group average performances achieved by the rats with the unocluded and occluded object views are reported, respectively, by the red curves and stars. (**c**) *L*_1_ distance between the pattern of accuracy of the rats across the whole set of unoccluded images (i.e., all red curves in **b**) and the patterns of accuracy achieved by the SVMs on the same images across VGG-16 layers. The shades of gray indicate the number of units that were sampled from each layer and fed to the SVMs (see colorbar on the right of **e**). The shaded areas are SEM across 5 classification runs. The histogram reports the occurrence of the minima of the *L*_1_ distance across classification runs and sizes of the populations fed to the SVMs. (**d**) Same as in (**c**), but with the *L*_1_ distance computed between the accuracy patterns of the rats and the SVMs in the case of the partially occluded images.(**e**) Pearson correlation coefficient between the accuracy patterns achieved by the rats and the SVMs across the entire stimulus set (i.e., pooling unoccluded and occluded views) as a function of VGG-16 depth. Shaded areas are SEM as in **c**. The histogram shows the distribution of the peaks of the correlation coefficient across classification runs and population sizes.

The test performed with VGG-16 followed the same procedure as the one described in the previous section, measuring the accuracy of linear SVMs to classify the two objects (presented across the full set of transformations used in ref.^5^), based on the activations of sub-populations of units sampled from each layer of the network. Also in this case, before being fed to the CNN, each image was first preprocessed through the augmentation pipeline. Given the randomness of the augmentation process and of the bubble masks, this resulted in two independent sets with thousands of images each - one to be used for training the SVM classifiers and another one to test their performances (see Methods).

The gray curves in Figure 4b show the test accuracies of the SVMs across the depth of the network (encoded by the shade of gray) as a function of the magnitude of the four transformations probed by Alemi-Neissi and colleagues^5^. Accuracy in the very initial layers (light gray) was close to chance, poorly modulated over the transformation axes, and even lower than the performance achieved by training an SVM to decode object identity directly from the pixel representations of the object views (cyan curves). It then progressively increased along the depth of the network, reaching, in middle layers (mid gray), an overall magnitude that was similar to the one measured for the rats (red curves). Concomitantly, the accuracy curves became increasingly closer to those of the rats also in terms of shape, eventually displaying a similar transformation dependence in middle layers. Accuracy kept increasing with the depth of the network, substantially surpassing rat performance and approaching 100% correct in the fully-connected layers (darkest gray). At the same time, the shape of the accuracy curves remained somewhat consistent with that observed for the rat curves even in deep convolutional layers.

The figure also reports a comparison of rat and SVM performances (red and gray stars, respectively) on the object views that, in the study of Alemi-Neissi and colleagues^5^, were partially occluded by the bubble masks. Rat recognition was quite robust to such manipulations, suffering only a minor drop of classification accuracy (the red stars are either at the same height or just below the corresponding dots on the red curves, reporting the performance on the unoccluded views). By contrast, the SVM classifiers, built over the representations provided by VGG layers, were afflicted by substantial performance losses (again, the accuracy yielded by the very initial layers was lower than that afforded by the pixel-level representation; compare light gray and cyan stars). Only the deepest layers (darkest stars) afforded performances that were comparable with those of the rats.

The trends reported in Figure 4b refer to experiments where 100K VGG units were sampled per layer, but these results were very robust across sub-population sizes and multiple runs of the classification procedure. This can be appreciated by inspecting the curves showing the L1 distances between the performance patterns observed for the rats and the SVM classifiers across VGG layers. In the case of the unoccluded object views, the smallest L1 distance was consistently reached in the middle of the network, with a sharp minimum in layer 8, regardless of population size (Fig. 4c). By contrast, for the partially occluded views, the L1 distance remained large and remarkably stable across the first half of the network, dropping sharply around layer 8 but then still decreasing till the very last layers (Fig. 4d). As a result, most L1 minimina were very concentrated in the final, fully connected layers.

As done in the previous section, we also computed the Pearson correlation coefficient between the accuracy patterns measured for the rats and the SVM classifiers across all tested image transformations (i.e., both unoccluded and occluded views). This similarity metric increased steadily till the middle layer, reaching a stable plateau in the second half of the network, with most minima concentrated between layers 9 and 12 (Fig. 4e).

Overall, comparing these results with those of the previous section further reaffirms the sophistication of rat object vision. In fact, when visual objects underwent a richer variety of image transformations, compared to those tested by Zoccolan and colleagues^4^, the pattern of rat discrimination performances could only be captured by deep convolutional layers of VGG-16. Most strikingly, the robustness of rat recognition to severe occlusion of the object views was only matched by the network in the final, fully connected layers.

### Comparing visual processing strategies in rats and CNNs

When using a convolutional neural network to model biological vision, comparing the two systems in terms of their discrimination accuracy only provides a first-order assessment of the model’s ability to capture the processing performed by the biological system. In fact, the same classification choices can in principle be supported by very different visual strategies, i.e., by very different sets of diagnostic features in the input images^35^. For instance, recent studies have shown that deep neural networks, despite their high classification accuracies, often employ image processing strategies that are substantially different from those used by humans^36^. This imposes a limit on the ability of neuronal network models to account for the tuning of ventral stream neurons - a limit that can be overcome by enforcing an alignment with human perceptual strategies^37,38^.

Inspired by these previous studies, we extended our assessment of rat object vision using VGG-16, by including a comparison at the level of visual processing strategies. This was possible thanks previous work of our group, in which two different classification image approaches have been applied to uncover the diagnostic features used by rats to discriminate visual objects^5,11,12^. These approaches are based on: 1) the already mentioned occlusion of object views by random bubble masks (see previous section and Fig. 4a)^5^; and 2) the presentation of randomly morphed variants of a previously learned target object^12^. By properly processing rat responses to these altered object conditions, both approaches allowed estimating the perceptual templates deployed by rats to recognize visual objects across various image transformations. In our tests with VGG-16, we adapted these procedures to infer the visual features underlying the choices of the SVM classifiers that were built over the representations provided by the network’s layers. This allowed comparing the rats and the CNN in terms of the similarity and generalization across transformations of their perceptual strategies.

### Rat image processing strategy is more view-invariant than VGG-16 strategy and more consistent with that of an ideal observer model

Alemi-Neissi and colleagues^5^ collected rat responses to partially occluded versions of previously learned visual objects (Fig. 4a). This allowed applying a classification image approach known as the Bubbles method, originally developed to infer the object classification strategies of humans^55^, and later applied also to other species, such as pigeons^56,57^, monkeys^58,59^ and, in the work of our group and others, rats^5,11,60^.

Intuitively, partial occlusion of visual objects impairs their processing, but the extent to which recognition becomes harder depends on what parts of an object are masked. If visual features that are critical to identify the object are erased, this will likely lead to mis-classification. Conversely, occluding image features that are not very diagnostic of object identity will not alter classification. As originally proposed by Gosselin and Schyns^55^, critical (or salient) features can be inferred by measuring the correlation between the random bubble masks and the behavioral responses of the observer to the corresponding, partially occluded object conditions (see Methods for details).

In our tests, we adapted this experimental procedure to CNNs by feeding VGG-16 with partially occluded object views and recording how the linear SVMs, trained on layer-specific activations, classified these images (see the *strategy inference* pipeline in the top branch of Fig. 1). As already done for the analysis shown in Fig. 4, the SVMs were trained using thousands of unmasked and randomly masked object views, equally divided in these two categories (see Methods). This was done to closely match the task that had been administered to the rats, who received feedback about the correctness of their responses to both clean and occluded images^5^. In addition, early tests revealed that inclusion of masked objects in the training pool was essential to avoid degenerate classification performances on the these conditions (i.e., perfect classification of one object and concomitant “perfect” mis-classification of the other one). This is consistent with the poor generalization afforded by VGG representations to masked images, as already shown in Fig. 4b. After training, each classifier was exposed to 3000 partially occluded images for each of the object views that had been tested with bubble masks in the rat experiment^5^ (i.e., the 7 views indicated by the stars in Fig. 4b plus the default view, yielding a total of 16 views, considering the two objects).

Figure 5 shows, for each of the 8 views of the “tripod” object, the group average saliency maps that were obtained for the rats by Alemi-Neissi and colleagues^5^ (2nd row), as well as those obtained for one example animal (3rd row). The figure also reports the saliency maps that we computed for the SVM classifiers trained on VGG units’ activations in a few convolutional layers (rows 4 to 7) and on the pixel representations (last row). Following our rat study^5^, the figure also shows the saliency maps obtained for an ideal-observer model (top row) - i.e., a classifier that had stored in memory, as templates, the eight views each object could take and that performed a template-matching operation (dot product) between each occluded input image and these 16 templates (assigning the object label based on the template that yielded the best match; see ref.^5^ for details). This simulated observer was “ideal” in the sense that had complete knowledge of the variability of object appearances that was present in the stimulus set and could thus solve the invariant task optimally. In Figure 5, the intensity of the maps is rendered in shades of grays, where light and dark pixels indicate, respectively, strong correlation and anticorrelation between the visibility of the pixels through the masks and correct classification of the object. That is, light and dark shades indicate portions of the object views that are, respectively, diagnostic (or salient) and anti-diagnostic (or anti-salient) with respect to object identity. The red and cyan patches highlight those object regions that were significantly salient and anti-salient according to a permutation test (*p <* 0.05; Methods).

**Figure 5:**
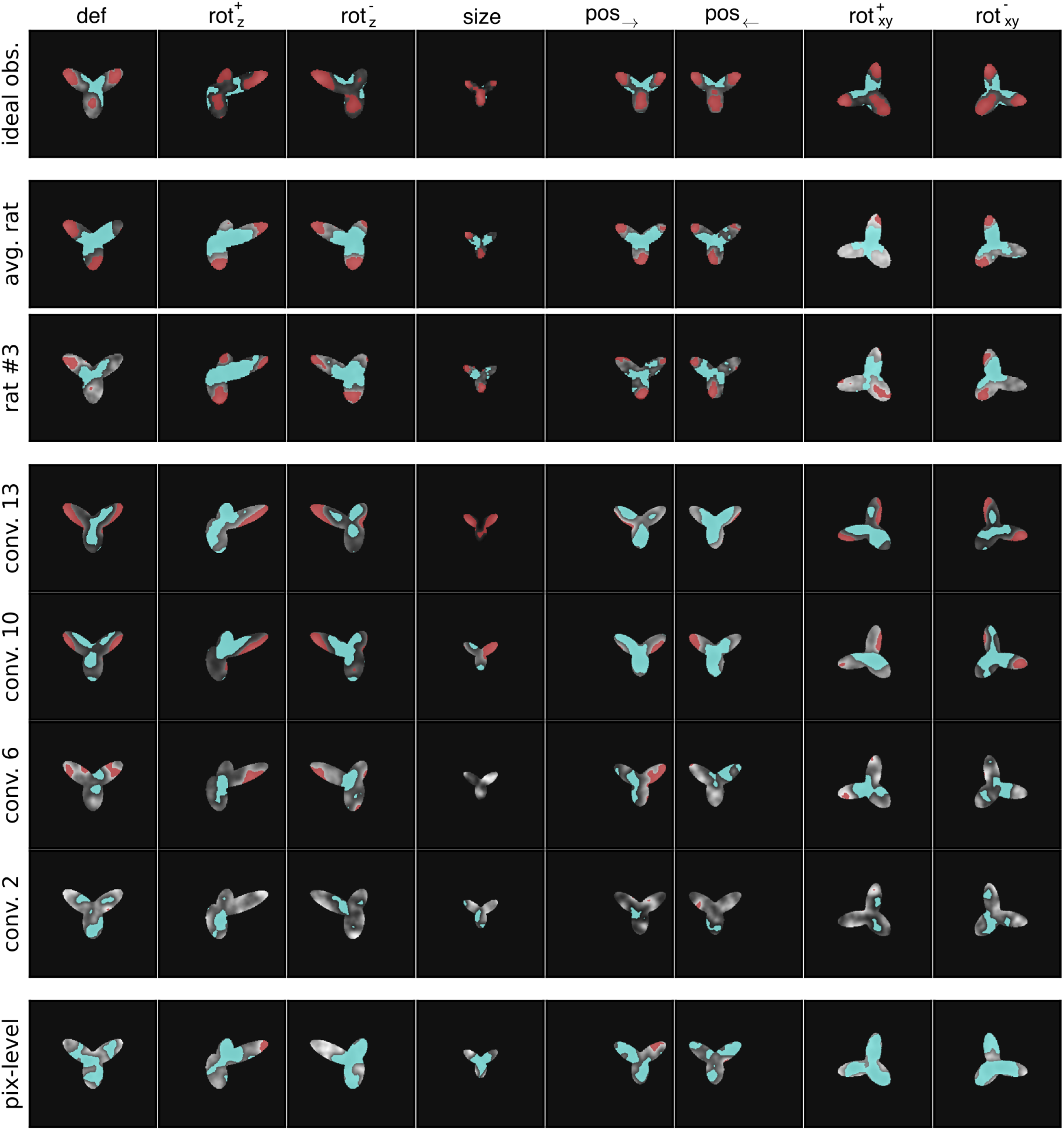
Comparing visual strategies among rats, VGG-16 layers, the pixel-based representation and an ideal observer model. The image classification approach known as the *Bubbles method* was applied to uncover the saliency maps used to recognize the *tripod* object by four different “perceptual”systems: 1) an ideal observer model (described in the main text; first row); 2) the rats, with the group average maps shown in the second row and the maps of an example animal (rat #3; see ref.^5^) shown in the third row; 3) linear SVMs applied to VGG-16 representations of the object in four checkpoint layers (rows four to seven); and 4) linear SVMs applied to the object’s pixel-based representation. In each map, the lightness of the gray scale indicates how well the visibility of a pixel correlated with the correct classification of the object (see Methods). The red and cyan colors indicates pixels that were, respectively, significantly correlated and anti-correlated with the correctness of the classification (*p <* 0.05; permutation test). These pixels form, respectively, significantly salient and anti-salient regions.

As already illustrated in ref.^5^, the saliency maps of both the average and the example rats featured multiple salient regions, located at the tips of the lobes of the tripod, as well as a central anti-salient area, located at the lobes’ intersection. Importantly, the relative position and size of the salient features was well preserved across views. By comparison, the saliency maps yielded by the representations in the pixel space and in VGG early layers were much less sharp, failing, in most views, to reach significance (i.e., little or no red patches are visible in the maps in the bottom rows of the figure). Also, the location of the salient regions (light gray) varied considerably from view to view. Stronger salient features emerged in late convolutional layers, although their shape was only partially consistent with that of the features in the rat maps, being located mostly at the edges rather than at the tips of the lobes. In general, regardless of the layer considered, the similarity between rat and VGG (or pixel) maps appeared to be low. By contrast, as already reported by Alemi-Neissi and colleagues^5^, both the average and individual rat saliency maps displayed a remarkable similarity with those of the ideal observer model.

These observations were quantified in Figure 6a, which reports the average Pearson correlation coefficient between the saliency maps obtained for the rats and those measured across VGG layers, with the average being computed across 6 rats and the 8 views of the tripod object (gray curve). This correlation remained very low across the whole depth of the network, being close to the one measured between the saliency maps of the rats and the maps obtained for the pixel-based representation (cyan dashed line). By comparison, the correlation of rat saliency maps with those obtained for the ideal observer was substantially higher (about 0.4; red dashed line), and an even larger value was observed when the average rat saliency maps were considered, as originally done by Alemi-Neissi and colleagues^5^ (about 0.55; green dashed line). Not surprisingly, the correlation of VGG saliency maps with those of the ideal observer was much lower (red solid curve) and not different from that with the maps yielded by the pixel representation (cyan solid curve). This indicates that rats were better than VGG-16 at discovering those portions of the “tripod” object that are more diagnostic of its identity in the face of view changes and partial occlusion. The network appeared to slighlty improve the optimality of its visual strategy across the convolutional layers (see the mild increase of the solid red curve), but never reached the same level of similarity with the strategy of the ideal observer as attained by the rats.

**Figure 6:**
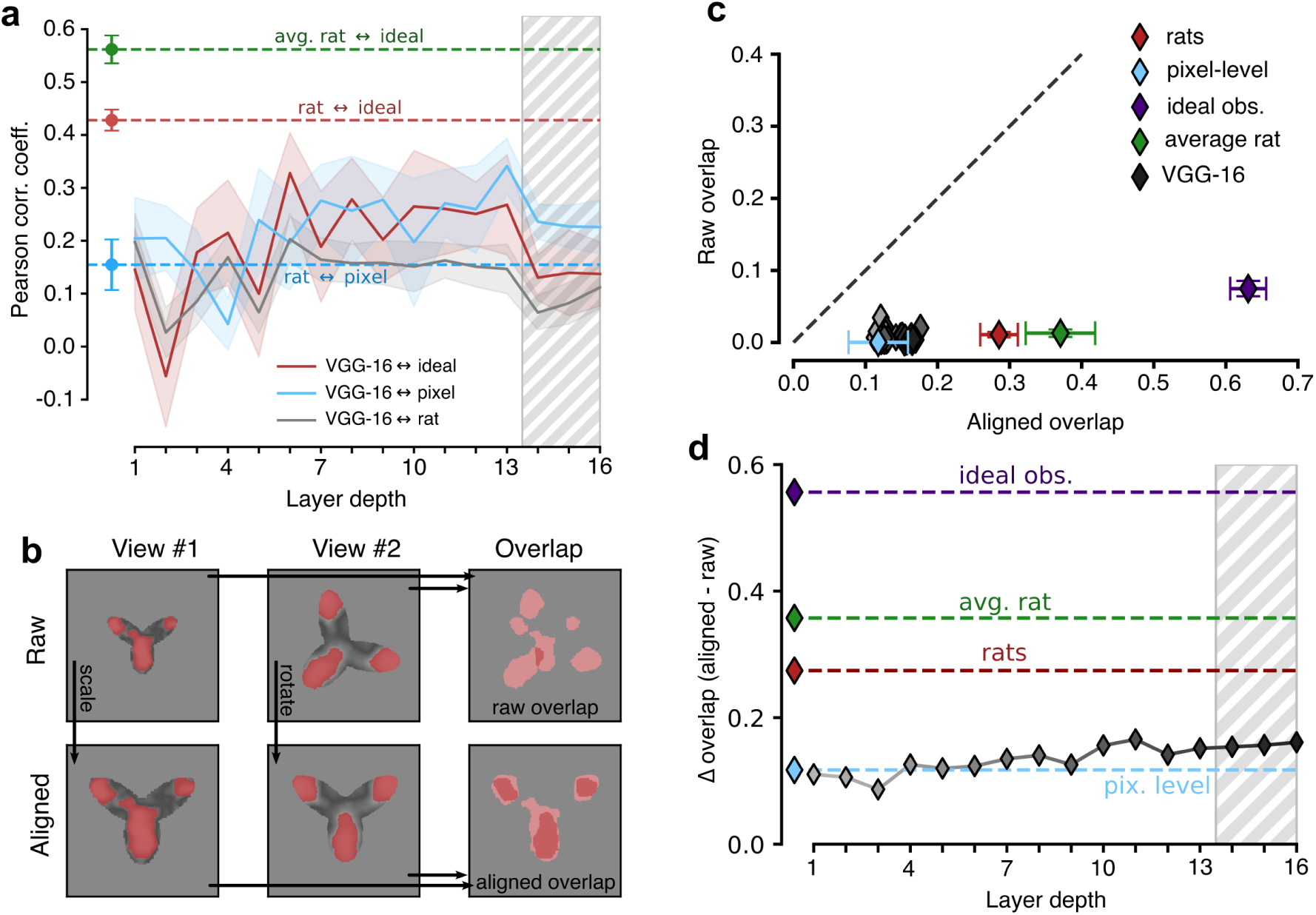
Superior view-invariance of rat visual perceptual strategies. (**a**) Pearson correlation coefficients between the saliency maps obtained for different pairs of “perceptual” systems: 1) rats vs. linear SVMs applied to VGG-16, layer-based representations (solid gray curve; the shaded area is the SEM across rats and object views); 2) rats vs. the ideal observer model (dashed red line; the error bar is the SEM across rats and object views); 3) average rat vs. the ideal observe model (dashed green line; the error bar is the SEM across object views); 4) rats vs. linear SVMs applied to the pixel-based representations (dashed cyan line; the error bar is the SEM across rats and object views); 5) SVMs applied to the VGG-16, layer-based vs. the pixel-based representations (solid cyan curve; the shaded area is the SEM across object views); and 6) SVMs applied to the VGG-16, layer-based representations vs. the ideal observer model (solid red curve; the shaded area is the SEM across object views). (**b**) Illustration of the procedure to compute the *raw* (top) and *aligned* (bottom) overlaps between the salient regions obtained for two different views of the tripod object (see main text). (**c**) Scatter plot reporting the *aligned* and *raw* overlaps of the salient regions obtained for the tripod object of the various “perceptual” systems under exam: 1) the rats (red); 2) the average rat (green); 3) the ideal observer model (purple); 4) the pixel-based representation (cyan); and 5) representations across VGG-16 layers, with the shades of gray (from light to dark) coding for the layer depth (from early to deep). Each point is the average over all the views of the tripods and all the rats (for the red point). Error bars are SEM (in the case of VGG-16 layers, SEM are not reported for clarity). (**d**) Difference between the *aligned* and *raw* overlap for the various “perceptual” systems shown in **c**.

This discrepancy between rat and VGG perceptual strategies could be interpreted in different ways. If one considers CNNs as the benchmark model systems for advanced visual processing, failure of rat saliency maps to tightly align with those of VGG would imply that rats process visual objects using lower-level, less refined strategies, as compared to the network. This conclusion would be in agreement with recent reports suggesting that rats rely on coarser contrast features than CNNs and primates do^61^. However, this interpretation would be at odd with the observation that rat perceptual strategies match those of the ideal observer model way better than VGG strategies do. Given that the ideal observer is, by construction, optimally view-invariant, this would suggest a superior ability of rats at discovering and deploying transformation-tolerant perceptual templates.

To test this hypothesis, we measured the invariance across object views of the visual strategies inferred for the rats, the VGG representations, the pixel representation and the ideal observer model. This property can be quantified by computing the extent to which the salient regions obtained for two different object views overlap. Critically, the overlap can be computed following two very different approaches, which allow distinguishing low-level, pixel-based image processing from invariant, feature-based recognition. As explained in ref.^5^ and illustrated in Figure 6b, given the saliency maps obtained for two views of an object, one can simply compute their “raw” overlap over the image plane (rightmost image in the top row). Otherwise, one can first reverse the transformations that produced the two views, realigning them back to the default view of the object (in the figure, this process is indicated by the arrows connecting the scaled and in-plane rotated views of the tripod, on the top, to their realigned versions, on the bottom), and then compute the resulting “aligned” overlap of the corresponding saliency maps (rightmost image in the bottom row).

The two overlap metrics have a very different meaning. A large raw overlap indicates that the observer consistently relies on the same portion of the visual display to recognize the object, thus suggesting the existence of some screen-locked, transformation-preserved cues that remain diagnostic of object identity despite view changes. This means that the supposedly invariant task can be trivially solved using a screen-centered, pixel-based strategy, which is easy for the observer to discover and apply. By contrast, a large aligned overlap means that the observer consistently extracts from the objects the same visual features despite view changes, where “same” refers here not to absolute screen coordinates but to the position, size and shape of the features relative to the structure of the object (e.g., the tips of the lobes of the tripod object). In other words, the aligned overlap measures the extent to which the visual strategy is object-aware and view-independent.

As shown by Alemi-Neissi and colleagues^5^, in the case of the rats, the aligned overlap was substantially larger than the raw overlap. This was particular evident for the tripod object (red diamond in Fig. 6c), for which the raw overlap (on average, across view pairs) was virtually zero, while the aligned overlap was close to 0.3 (i.e., about 30% of the union of the saliency maps obtained for two views overlapped, on average). Based on this analysis, Alemi-Neissi and colleagues^5^ concluded that rat recognition could not be accounted for by a low-level, screen-centered strategy and relied instead on object-centered visual features that were fairly well preserved across transformations. Figure 6c further reinforces this conclusion by showing that, when the idiosyncratic strategies of the individual rats were combined in the group average saliency maps (i.e., those shown in the second row of Fig. 5), the aligned overlap became even larger (green diamond), approaching 0.4. This value was not too far from that obtained for the ideal observer model (about 0.6; purple diamond), which, not surprisingly, displayed the highest, “non trivial” view-invariance.

In our study, we replicated this analysis for the saliency maps obtained for the SVM classifiers trained on the pixel representation and on VGG layers’ activations. The aligned overlap obtained for the pixel-based maps (cyan diamond) was just one third of the one observed for the individual rats and about one forth of that measured for the average rat. The aligned overlaps computed for the VGG maps (grey-shaded diamonds, with the depth of the network increasing from light to dark shades) were also very low in the initial layers, and close to the overlap obtained for the pixel-based maps. Interestingly, despite a tendency to increase in deeper layers, the aligned overlaps remained substantially lower than the overlap measured for the rats, never surpassing 0.2.

These trends were further quantified by plotting the difference between the aligned and raw overlap as a function of the network depth (Fig. 6d; gray curve). Although a modest increase could be observed, the network never reached the level measured for the individual rats (dashed red line) or for the average rat (dashed green line). The difference remained very close to the one observed for the pixel-based maps (cyan dashed line) and very distant from the one measured for the ideal observer (purple dashed line).

Overall, this analysis shows that, along the continuum that goes from the low-level, poorly invariant strategy of the pixel-based representation to the highly invariant perceptual templates of the ideal observer, the visual strategies afforded by VGG representations sit very close to the pixel one. Rat perceptual templates substantially depart from such poorly invariant strategies and tend towards the maximally invariant templates attained by the ideal observer. The implications of this finding are examined in the Discussion, in the context of previous work comparing image processing in humans and CNNs.

### Rat perception is more invariant than VGG-16 representations to reduction of visual objects to their outlines

An alternative classification image approach to infer the perceptual strategies underlying visual object recognition in rats has been developed by Djurdevic and colleagues^12^. In that study, the animals were initially trained to discriminate the tripod object from a set of *distractor* objects (shown in ref.^12^). Since the image of the tripod was rendered from a 3D model, its appearance could be parametrically altered by random variations of its structural parts, resulting in a new set of stimuli, referred to as “random tripods” (see examples in Fig. 7a). Once a rat had learned to successfully discriminate the tripod from the distractors, it started to be presented also with the random tripods. The rats spontaneously classified these stimuli as belonging to either the tripod or the distractor category, based on how similar the objects were perceived to the original tripod. Importantly, the animals never received any feedback about the correctness of their choices. This allowed testing their pure, spontaneous generalization to a new set of previously unseen images. In addition, the experimental design of ref.^12^ allowed probing the invariance of the generalization process, since the animals were also tested with scaled and outline versions of the random tripods (an example of the latter is shown in Fig. 7b, left).

**Figure 7:**
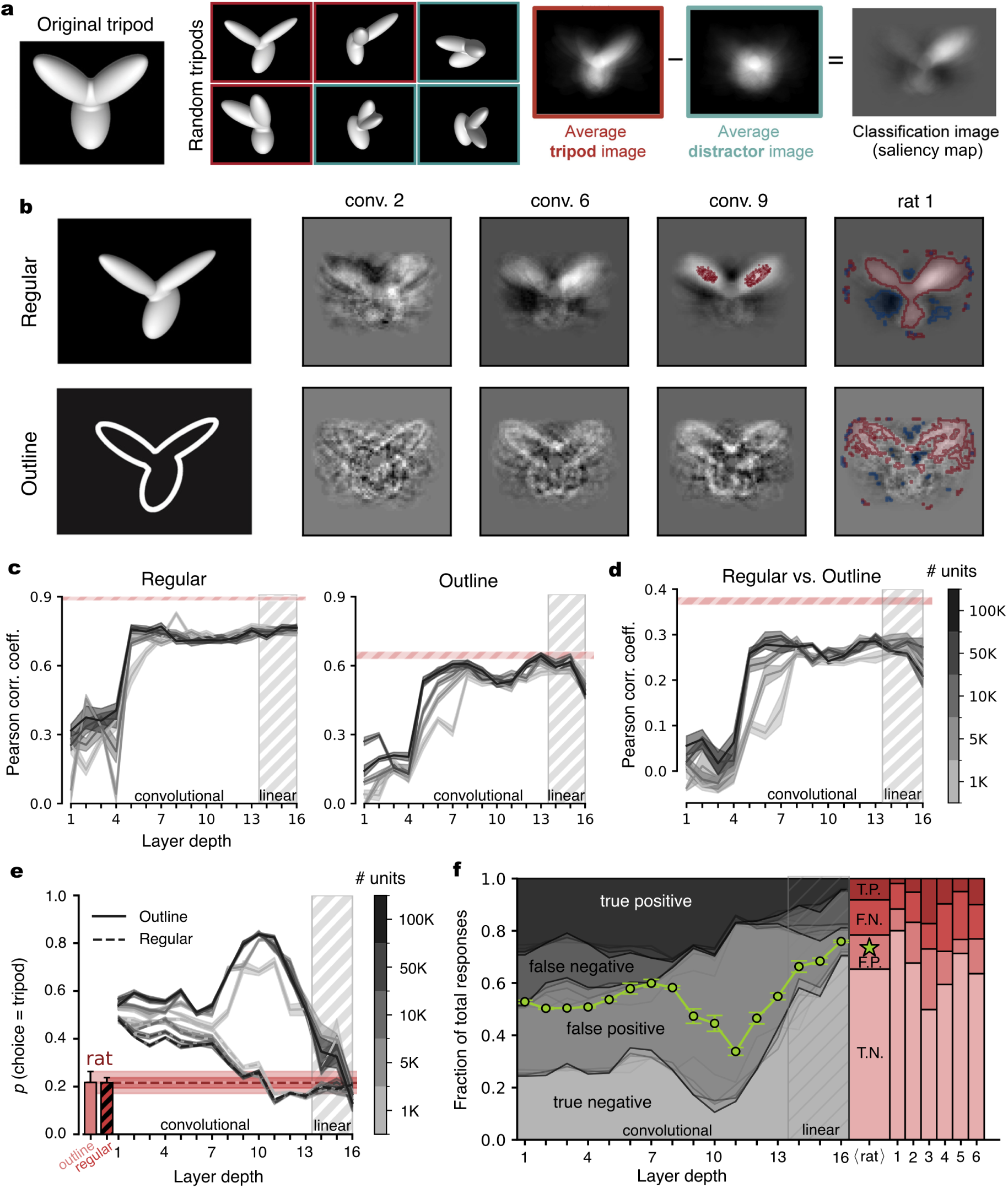
Superior generalization of rat perception to outline versions of visual objects. (**a**) Stimulus set and experimental design originally used by Djurdevic and collegaues^12^. Subjects were trained to recognize the tripod (left) from a set of distractor objects (not shown). They were then tested against random structural variations of the tripods. A saliency map (rightmost image) could thus be obtained by averaging the random tripods that were classified as the tripod (red frames) and subtracting the average of those that were classified as distractors (cyan frames). The same procedure was applied in our experiments with VGG-16. (**b**) The saliency maps obtained for linear SVMs applied to the representations in three convolutional layers of VGG-16are shown alongside the saliency maps of an example rat from ref.^12^. The maps were obtained for two classes of stimuli: regular (i.e., full-body) objects and their outline versions (examples shown on the left). In the maps, light- and dark-gray regions refer to pixels that were, respectively, correlated and anti-correlated with reporting the “tripod” choice. The red and cyan contours mark regions for which such correlations and anti-correlations were statistically significant (*p <* 0.05 on permutation test with 100 repetitions). (c) Pearson correlation coefficient between the saliency maps obtained for the rats and those computed for linear SVMs applied to VGG-16, layer-based representations using either regular (left) or outline (right) random tripods (solid gray curves; the shaded areas are SEMs across rats and classification runs). The shades of gray indicate the number of units that were sampled from each layer and fed to the SVMs (see colorbar in e). The shaded red stripes mark the SEMs (centered on the averages) of the correlation coefficients between the saliency maps of the 6 rats tested by Djurdevic and colleagues^12^. (d) Pearson correlation coefficient between the saliency maps obtained for linear SVMs applied to VGG-16, layer-based representations in the case of matching views of the regular and outline random tripods (the shaded areas are SEMs across classification runs). The shaded red stripe indicates the same analysis for the saliency maps obtained for the rats (SEM centered on the rat group average). (e) Fraction of random tripods that were classified as the *tripod*, in the case of regular and outline objects (dashed and solid lines, respectively), by the rats (red bars/lines) and by linear SVMs applied to VGG-16, layer-based representations (gray curves). Shades areas and shades of gray as in d. (f) Visualization of the confusion matrix obtained when considering the classification of the regular random tripods performed by the rats (red bars) or by the SVMs applied to VGG-16, layer-based representations (gray curves) as the ground truth, against which to compare the classification of the outline versions of the stimuli. The shade of gray of the curves indicates the the number of units that were sampled from each layer and fed to the SVMs (see colorbar in f). The green dots and the star show the classification accuracy achieved, respectively, by the SVM classifiers and by the rats (group average).

As in the study of Alemi-Neissi and colleagues^5^, the ultimate goal of this experiment was to uncover the perceptual templates used by the rats to recognize the tripod object across transformations. To this aim, a saliency map was obtained from the pattern of responses of each animal to the random tripods, by computing the difference between the average of the images that had been classified as the tripod and the average of those that had been classified as a distractor (see Fig. 7a and the Methods). Compared to the Bubbles method used in ref.^5^, this approach had the advantage that it did not require masking the objects. This, in turn, allowed looking for salient an anti-salient features also outside the boundaries of the tripod. In addition, the resulting saliency maps were obtained from object conditions (the random tripods) that were better matched to the originally learned tripod in terms of both low-level (e.g., luminosity) and higher-order (overall geometry) properties.

Djurdevic and colleagues^12^ tested in this experiment six rats, finding classification images with salient regions typically encompassing two or three lobes of the tripod and anti-salient regions located at the lobes’ intersections (see the red and cyan regions in the map shown for an example rat in Fig. 7b, last column). Small differences existed among the maps inferred for different animals, which were impactful enough to account for the different performances of the animals with the distractor objects. In addition, the saliency maps obtained for a rat from the regular, small, and outline versions of the random tripods were highly consistent, highlighting, once more, the invariance of rat recognition strategy.

In our study, we replicated the original experiment of Djurdevic and colleagues^12^ in an artificial setting. We trained a linear SVM for each layer of VGG-16 to correctly discriminate the original tripod from the pool of distractor objects using the inner representation afforded by a sub-population of units in each layer (as in our previous tests, sub-populations of different sizes were used). We then fed the network with the set of random tripods and we recorded the classification labels assigned by the SVMs to these stimuli, in a pure generalization setting that matched the test applied to the rats. Following an analogous procedure to the one presented in ref.^12^, we computed, for each layer of the CNN, a saliency map that revealed which visual features were critical for successful discrimination of the tripod from the distractor objects.

Example saliency maps for the regular (i.e., full body) and outline conditions are reported in Figure 7b for three convolutional layers of VGG-16, alongside the saliency maps of an example rat. From visual inspection, the saliency maps obtained from VGG representations seemed to progressively converge to the map measured for rats. To quantify this intuition, we computed the average Pearson Correlation coefficient between VGG and rats’ saliency maps as a function of the depth of the network (Fig. 7c, gray lines). We also computed the average Pearson Correlation coefficient among the maps of the six rats, to serve as a consistency benchmark (the red shaded area in the figure indicates the the region encompassed by *±*SD over such average, rat-wise correlation).

For the maps extracted from the responses to the regular random tripods (left plot), the correlation started low and then increased sharply from layer 4 to layer 5, reaching a plateau that was remarkably stable over the rest of the network - a finding that was robust across the different population scales we explored. Differently from what observed in the comparison with the study of Alemi-Neissi and colleagues^5^ (see Fig. 6a), this asymptotic correlation was high (around 0.7), thus showing that middle to late VGG layers yielded discriminatory visual features that were quite consistent with those of the rats. At the same time, such consistency did not reach the one among rat maps, whose correlation was close to 0.9. This indicates that, even in deeper layers, VGG saliency maps did not fully capture the perceptual strategies deployed by the rats. The correlation between VGG and rats’ saliency maps followed a similar trend also in the case of the outline stimuli (Fig. 7c, right). In this case, however, the maximal correlation matched the one measured among the rats. In addition, following an initial plateau in the middle of the network, the correlation slightly decreased to further grow in the final convolutional layers.

As already mentioned, Djurdevic and colleagues^12^ found that the saliency maps obtained from the various, transformed versions of the random tripods were quite consistent - even in the case of radical transformations such as the change from regular to outline stimuli. Such consistency is quantified in Figure 7d, which reports the within-rat correlation between the saliency maps yielded by regular and outline random tripods, averaged across the six animals (red shaded stripe). The magnitude of the correlation approached 0.4, which is no small value considering how different the stimuli yielding the two kinds of saliency maps were and, as a result, how scattered the outline-based saliency maps were, as compared to the regular-based maps (contrast the two images in the last column of Fig. 7b). The same correlation metric was computed between the maps obtained from regular and outline stimuli for each layer of VGG-16 (gray curves). The correlation sharply increased between layer 5 and 7 to reach a relatively stable plateau, but failed to match the value measured for the rats, even in deep layers. This confirmed the conclusion of the previous section - i.e., that rats employ perceptual strategies that are more invariant to challenging image transformations than those afforded by a fully-trained CNN. As a final step in our analysis, we took inspiration from one of the results presented in the previous sections - i.e., the poor generalization of VGG representations to the partially occluded object conditions (star symbols in Fig. 4b). This finding suggests that, despite its proficiency with challenging image sets like Imagenet, VGG-16 is quite sensitive to severe image manipulations - possibly, those that drastically alter the surface area of the objects, strongly reducing their luminosity, thus yielding out-of-distribution samples. The stimulus set of ref.^12^ offered the possibility to further investigate this phenomenon, given that transforming the random tripods into their outlines produced image changes as severe as the application of the bubble masks. Quite impressively, for the rats, these changes did not alter at all the average likelihood of classifying a random tripod as being the tripod, as opposed to belong to the distractors’ category. As shown by Djurdevic and colleagues^12^, the probability that a rat classified a random tripod as being the tripod was equally low (about 0.2) and virtually identical for both the regular and the outline stimuli (compare the red to the striped bar in Fig. 7e). This suggests that rats developed perceptual templates that were sharply tuned to the shape of the tripod and such templates were fully invariant to the transformation that changed the regular, full-body stimuli into their outline counterparts.

When we applied this analysis to the choices of the SVM classifiers trained on the tripod vs. distractor task based on VGG representations, we found several important differences with rat behavior. Up to the middle of the network, the probability to classify a random tripod as being the tripod (gray lines in Fig. 7e) was similar for the regular and outline stimuli, but much higher (around 0.5) than the one measured for the rats. This indicates that reading the activations of VGG units in early to middle layers did not allow the linear classifiers to find regions, within the representational spaces, that were specific enough for the shape of the tripod to not include most of its random, structural variations. Interestingly, starting from layer 8, the proportions of tripod choices for the outline and regular stimuli diverged, reaching, in layers 10-11, extremely large ( 0.8) and low (*<* 0.2) values, respectively. This indicates a complete lack of invariance in late convolutional layers to the outline transformation. Only in the very final convolutional and fully-connected layers the probability of tripod responses to the outline stimuli dropped, while that to the regular stimuli slightly increased, with both probabilities eventually converging to those measured for the rats.

To further investigate these trends, we also measured how consistent the classification of every random tripod was, when presented in its regular and outline version. While no correct label existed for the classification of the random tripods, one can quantify the consistency between the responses to the two variants of these stimuli (regular and outline) by considering the decisions for a given variant as the “ground truth” and then checking whether the choices for the other variant are consistent with these “true” labels. In our analysis, we used the classification of the regular random tripods to define their “correct” labels and we tested whether the classification of the same random tripods, but in their outline version, was in agreement with these labels. This yielded the confusion matrices shown in Figure 7f for VGG layers (gray shaded areas) and for the rats (red shaded bars). For the latter, despite some variability across animals, the classification accuracy (i.e., the consistency between responses to regular and outline random tripods) was high, with the sum of true negatives (i.e., random tripods classified as distractors in both their regular and outline variants) and true positives (i.e., random tripods classified as the tripod in both their regular and outline variants) being close to 70% on average across rats (green star). In the case of the network, classification accuracy was substantially lower till the very final layers (green line), mainly because of the large rate of false positives and the concomitantly low rate of true negatives, especially in layers 9-11 (in agreement with the strong divergence of tripod choices for regular and outline stimuli illustrated in Fig. 7e). Only the deepest, fully connected layers of the network displayed a confusion matrix that closely matched the one of the average rat. This further reinforces the conclusion that the full extent of the CNN is needed to model the invariance of rat object vision under the most challenging, out-of-distribution image manipulations.

## Discussion

The goal of our study was to evaluate the processing power of rat object vision by carrying out a comparison with a popular CNN: VGG-16. Our approach was similar to the one adopted in previous CNN-based assessments of the proficiency of humans and rats to recognize visual objects^27,28^. We estimated how hard it was for a rat to succeed in a given object recognition task by interrogating object representations across VGG layers to test how well they supported the task (*accuracy measurement* pipeline in Fig. 1). The depth of the layer yielding the best match with the pattern of rat discrimination accuracies was taken as a measure of how advanced the processing carried out by the animals to succeed in the task was. However, differently from Vinken and Op De Beeck^28^, our approach included: 1) a preprocessing/augmentation stage that closely reproduced the blurring, noise level and image variability experienced by the animals in the tasks (Fig. 2); 2) a focus on comparing how similarly recognition accuracy was modulated across object views in rats and CNN layers, rather than a comparison of absolute discrimination performances; and 3) a comparison between the “perceptual” strategies used by rats and CNNs to solve the same object recognition task (*strategy inference* pipeline in Fig. 1). Our experiments yielded three main results.

First, we found that about half the depth of VGG-16 is required to match the patterns of rat classification accuracy in tasks where objects underwent changes in position, size, in-depth and in-plane rotation (Figs. 3e-f and 4b-c). This conclusion is in disagreement with the study of Vinken and Op De Beeck^28^, who found that the very first layer of VGG-16 and Alexnet (an older, shallower CNN) was sufficient to achieve nearly perfect discrimination accuracy (i.e., well beyond that attained by the rats) in the object recognition task reported in ref.^4^. Such discrepancy is not surprising, given the underestimation of the actual image-level variability, noise and blurring experienced by the rats in the analysis of ref.^28^, as well as their focus on comparing absolute discrimination accuracies. On the other hand, Vinken and Op De Beeck^28^ also tested the extent to which the representations in CNN layers were able to account for the the pattern of rat behavioral accuracies over the tested object views, as reported in ref.^4^ (i.e., the matrix of performances shown in Fig. 3b). The rationale of this analysis is more consistent with that of our approach and, not surprisingly, the tests of Vinken and colleagues showed that mid to high-level convolutional layers were necessary to account for the modulation of rat performance across object conditions.

The second major finding of our study is that the entire depth of VGG-16, up to the final, fully connected layers, was required to account for rat ability to generalize to more radical image manipulations of visual objects: heavy occlusion and reduction of an object to its outline (Fig. 4d-e and Fig. 7e-f). In addition, generalization to occluded images was not achievable without including them in the training diet of the SVMs, so as to avoid degenerate performances on these stimuli. By comparison, the robustness of rat perception to these transformations (Fig. 4b and 7e-f) indicates that, when visual patterns are incomplete and undergo severe variations of luminosity, the relatively shallow rat visual cortical hierarchy^7,15,62^ displays all its generalization power - well beyond what revealed when the animals are tested with spatial transformations and rotations or even with naturalistic movies^28^, and well beyond what achieved by VGG-16 early to mid-level convolutional layers.

It could be argued that this ability lies in the pruning of low-level visual properties (such as luminance and contrast) that is carried out by the rat object-processing pathway^7^. However, we recently reported evidence for a similarly strong pruning taking place along VGG-16 early layers^54^. Thus, it is more likely that rat robustness to occlusion resides on processing mechanisms that are not purely feedforward, in agreement with the conclusions of previous studies suggesting a pivot role of recurrent computations in supporting pattern completion^63–67^. Interestingly, some of these studies showed that CNNs are not as tolerant as the human visual system to partial occlusion, but their robustness to this manipulation was substantially increased by the addition of recurrent connectivity^64,67^. Since a recurrent neural network can be unfolded, by adding extra layers for each recurrent step, to create a computationally equivalent feedforward network^68^, this can explain while the full depth of VGG-16 is necessary to match what rat visual cortex achieves with way less (but highly recurrent) processing stages - i.e., 6-7 visual areas from retina to the highest cortical areas of the object processing pathway^7,15,62^.

As for generalization to outlines, the struggle of VGG-16 to deal with such image manipulation is fully consistent with previous studies showing how convolutional networks are way less proficient than humans in recognizing objects that are deprived of texture, such us silhouettes and outlines (with the latter being particularly challenging for CNNs)^69–71^. This indicate that CNNs process images by exploiting sets of features that are substantially different from those used by humans - i.e., without extracting a representation of the global shape of visual objects^39,70^, but heavily relying instead on texture information^71^. By contrast, rat proficiency in dealing with outlines indicates an intriguing similarity in the way rats and humans process visual shapes.

Taken together, these considerations suggest that the merit of the layer-by-layer comparison between CNNs’ representations and rat object vision is in the possibility to bridge rat and human visual perception. Which depth of a CNN yields the best match with rat discrimination accuracy does not matter much per se. It is not surprising that CNNs largely surpass rats in terms of performance magnitude in object recognition tasks - they do so with human performance too^72^. The more intriguing question is whether the way visual perception departs from the processing carried out by CNNs is similar in rats and humans. Our findings indicate that this is indeed the case - and to a surprising extent. Not only humans and rats display similarly high robustness to heavy image occlusion and reduction of objects to outlines - a robustness that CNNs struggle to achieve. Eberhardt and colleagues^27^ have shown that, when humans are tested in a speeded image classification task, their accuracy is surpassed by the networks of the VGG family already in early to mid-level layers, while the correlation between human and CNN confidence in classifying the images peaks in mid to late-level layers.

The consistency of these findings with those of our experiments (Figs. Fig. 3e-f and 4b-c) suggests that the core mechanisms underlying shape processing in humans and rats are similar - and are similarly different from those of CNNs. In the spirit of what suggested for humans^68,73^, rats, under the pressure of the discrimination task and of the water restriction regime, likely process incoming visual objects via a fast feedforward sweep through the visual hierarchy. This allows them to achieve performances that are high enough (even in the face of translation, rotation and scaling) to fulfill the goal of obtaining the liquid reward as fast as possible (note that achieving very high accuracy on individual stimulus presentations at the cost of a slower response time and, therefore, of a lower number of stimulus presentations, may not be optimal in terms of the overall amount of reward achieved over the course of the experiment). However, the animals, as postulated for humans, when facing way more challenging image manipulations (such as occlusion and outlines), may flexibly increase the depth of visual processing by engaging recurrent processes^67,68,73,74^ - a strategy that in standard CNNs would require exploiting the full depth of the purely feedforward architecture.

The existence of systematic differences in the processing of shape information between CNNs and biological brains was confirmed by the comparison we carried out at the level of perceptual strategies - the third main finding of our study. As already shown by Alemi-Neissi and colleagues^5^, rat perceptual strategy, as revealed by the animals’ responses to partially occluded images, was quite consistent with that of an ideal observer, which, by design, was optimally tolerant to object transformations (Fig. 5 and 6a). By contrast, rat strategy correlated poorly with the saliency maps extracted across VGG layers, which, not surprisingly, were also poorly consistent with those of the ideal observer (Fig. 6a). More importantly, we found that the diagnostic features used by rats to recognize visual objects across view changes were considerably more invariant than those afforded by the representations of the same objects along VGG layers, even the deepest ones (Fig. 6c-d) - a conclusion that was confirmed by comparing the saliency maps yielded by radically different appearances of visual objects, such as full-bodies vs outlines (Fig. 7d).

These findings, taken together with the results discussed in the previous paragraphs, suggest the existence of a trade-off between the discrimination accuracy a given representation can afford and the invariance of the set of visual features encoded by the representation, upon which the discrimination is based. VGG-16 deep layers support much higher classification accuracies of the unoccluded stimuli, as compared to rat performance. However, these larger accuracies are attained by relying on object representations that are more view-specific than the perceptual templates exploited by the rats. This seems consistent with the ability of deep CNNs to achieve virtually perfect training accuracy in classification tasks where images are randomly associated to arbitrary category labels^75^ - an impressive feat that, however, points to a fundamental difference with biological vision in terms of the ability to learn abstract category information^39,68^. Under this scenario, it is unclear which of the two systems actually encode objects in the most “advanced” way. Despite the larger accuracy it affords, the stronger view dependency of VGG-16 representations could be at the root of their lower ability to generalize to severe, out-of-distribution, image-level manipulations (such as occlusion and reduction to outlines).

Interestingly, these conclusions resonate with those of recent studies showing that the object recognition strategies deployed by deep nets are poorly aligned with those used by humans observers and with the saliency maps describing the tuning of IT neurons^36–38^. Moreover, such discrepancy has worsened as the discrimination accuracy on Imagenet of more recent (and larger) architectures has increased. This has led some authors to design a training routine to explicitly align the visual strategies of deep nets to those employed by humans, with the goal of obtaining more biologically relevant models of visual perception and cortical processing^37,38^. The importance of matching perceptual strategies between biological visual systems and neural networks is also highlighted by another study^76^, where a generative model of visual faces was used to parametrically control various properties of face features. The authors tested different neural network models in terms of their ability to predict human face similarity judgements, showing that the best models shared with humans a consistent set of diagnostic face landmarks. Yet other studies have tackled the same issue by complementing the training diet of CNNs with the addition of images where texture had been altered in such a way be uninformative about category labels^71^, or where blurring had been added to mimic the low resolution of peripheral retina^77^. As a result of this augmentation of the visual diet, CNNs acquired a stronger sensitivity to the shape of visual objects, as opposed to their texture, thus matching more closely human perception and also improving in terms of robustness to various forms of noise and distortions.

In the light of these considerations, the results of our study not only indicate a surprising degree of consistency between species so evolutionary and ecologically distant as rodents and primates, but also reinforce the need to further increase the robustness, generalizability, and biological consistency of CNNs^35,39^. For instance, in the spirit of refs.^71,77^, our study suggests that images with a variable amount of occlusion (e.g., via bubbles masks) should be included in the training diet of CNNs, to make them more robust to such manipulation. Also, in the spirit of refs.^37,38^, a high degree of consistency between the visual features used to recognize augmented versions of the same objects could be enforced in the training routine of CNNs, with the goal of learning more invariant shape representations of whole objects. Finally, in the spirit of refs.^67,68,74,78^, our study suggests that some amount of recurrent connectivity should be included in CNN architectures, to allow adjusting the depth of visual processing by flexibly selecting the number of repeated sweeps through the processing hierarchy, depending on the needs of the task at hand.

In summary, our work not only reasserts the sophistication of rat vision and, therefore, the importance of rodent models to dissect the neuronal circuits underlying visual perception. It also points to potentially interesting adjustments to the architecture and visual diet of CNNs to make their representations, robustness and generalization power closer to their biological counterparts.

## Methods

### Computation of the contrast sensitivity function

The contrast-sensitivity function (CSF; see Fig. 2a) provides a concise description of the acuity of an organism’s visual system, by assessing the contrast visibility threshold of static sine-wave gratings as a function of the stimulus’ spatial frequency. Operationally, the CSF can be estimated by administering a visual detection task, where an observer is required to report whether sine-wave gratings of variable spatial frequencies and contrast are detectable over a gray background. By letting the contrast *c* vary from full-visibility (i.e., the condition in which the difference between maximal and minimal luminance is the greatest) to no-visibility (i.e., uniform gray display), one can reconstruct the psychometric response functions *p* (detection*, c*; *ν*), where *p* is the probability of detecting the grating as a function of its contrast *c*, given a spatial frequency *ν* (see Fig. 2c). The contrast sensitivity threshold *ξ* (*ν*) can be defined in different ways^30,79,80^. The simplest approach is to set it to the contrast value yielding a criterion detection performance and is often chosen as the point of subjective equality - i.e., *ξ* such that *p* (detection*, ξ*; *ν*) = 50% (dashed red line in Fig. 2c). The contrast sensitivity *σ* (*ν*) (at a specific spatial frequency) is then computed as the reciprocal of the measured sensitivity threshold, i.e., *σ* (*ν*) = 1*/ξ* (*ν*). The CSF is thus estimated by measuring *σ* (*ν*) for different values of the spatial frequency *ν* of the gratings. This approach was used to compute the CSFs of the simulated observer shown in Fig. 2a (purple curves), as explained in the Results.

### Image augmentation pipeline

The image augmentation pipeline simulates the additional image variability produced by the head movements of the rats that, in our experiments, were not head fixed, but only slightly body restrained. In fact, the animals had the insert their head through a viewing whole, in order to face the stimulus display and get access to the array of touch sensors that allowed them to trigger stimulus presentation and report stimulus identity^4,5,12^. This ensured precise control over the viewing distance *d* (which, in all our experiments, was set to *d* = 30cm), but allowed for variability in head rotation around the three Euler angles (yaw, pitch, and roll) at the time when the rat sampled the visual stimulus. The images were shown on a monitor of size 48cm *×* 27cm (width *×* height), resulting in a size of 35*^◦^* of visual angle for the default views of the visual objects.

To estimate the impact of head rotation on the appearance of the visual objects used in our experiments, we leveraged previous work from our lab^32^, in which we characterized the variability of head orientation about the three Euler angles (for rats tested under equivalent experimental conditions as those used in refs.^4,5,12^) by means of head-tracking. Specifically, our previous study estimated the following ranges of variations of head angles: 60*^◦^*, 35*^◦^* and 20*^◦^* for pitch, roll and yaw respectively. This resulted in vertical translations (pitch), horizontal translations (yaw) and in-plane rotations (roll), the extent of which we quantified based on the geometry of the experimental setup.

The magnitude of the in-plane rotations directly corresponds to the magnitude of the roll rotations *θ_r_*. Therefore, in our augmentation pipeline, we assigned to any visual object fed to VGG-16 a random in-plane rotation *α* = *θ_r_*, with *θ_r_* sampled uniformly in *∼* [*−*17.5*^◦^,* +17.5*^◦^*]. To quantify the horizontal and vertical displacements 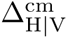 of object images due to variations in pitch *θ_p_* and yaw *θ_y_*, we computed

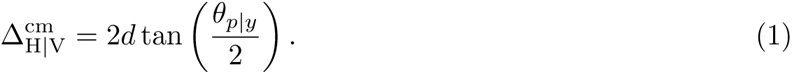

The resulting 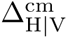 is expressed in cm. However, for our image augmentation pipeline, we used the PyTorch transforms.RandomAffine routine, which expects the *x*-*y* translations to be specified as a fraction of the original image size. We thus made a conversion from cm to pixel-count by knowing 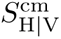, the size of the images presented to the rats in cm, and 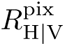, the screen resolution in pixels, and by computing the cm-to-pixel conversion factor as: 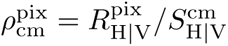. One can then express both the image size and the displacement in pixels as, respectively: 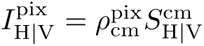 and 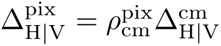. The *x*-*y* translations were then uniformly sampled as 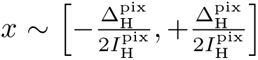 and 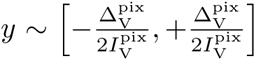. The full augmentation pipeline encompassed all the steps described above (see examples in Fig. 2e) and included also the addition of Gaussian blur and random Gaussian noise, as explained in the Results (see Fig. 2).

### Training and testing procedure of the linear SVMs on the object representations yielded by VGG-16 layers

In all our experiments, we used a VGG-16 network (from PyTorch) pre-trained on the ImageNet dataset. For each convolutional and fully-connected layer of the network, we built a Linear SVM classifier (using the sklearn.svm Python implementation) that was trained to predict the identity (i.e., the label) of the visual objects used in the rat experiments, using the activations of a subpopulation of units measured before the nonlinear ReLu gate. We measured these activations for all the units in a layer, following the presentation of the (augmented) stimulus conditions of a given rat experiment, and then we sampled a random sub-population with a given size. We explored the following range of population sizes: {10^3^, 5 *×* 10^3^, 10^4^, 5 *×* 10^4^, 10^5^}. Scaling beyond this value was limited due to memory constraints. However, not all VGG-16 layers have a total unit count greater than the biggest scale considered (for instance, the output layer only has 10^3^ units). In case the total number of units in a layer was smaller than the considered scale, we simply took the whole population of units. This only occurred for the linear, fully connected layers, as the last convolutional layer in VGG-16 has more than 10^5^ units.

Before being fed to the linear classifier, the sampled activations were independently normalized to have zero mean and unit variance across the set of stimulus conditions used for training (this was done with the scaler implementation from sklearn.preprocessing.StandardScaler). Thanks to the augmentation pipeline explained in the previous section, we could scale the number of both training and testing conditions beyond the original sets described in the rat studies, simply by re-sampling from each set and exploiting the additional variability introduced by the random augmentations. It should be noted that this procedure was adopted to only to achieve a more robust training/testing of the classifiers, but also because it more closely reflected the actual amount of image-level variability experienced by the rats.

In all experiments, we trained the linear classifiers on 5 *×* 10^3^ images sampled from the appropriate stimulus set, while the number of testing conditions varied based on the rat study under exam. In the experiments where the stimulus set of Zoccolan and colleagues^4^ was used, we tested the SVMs on 5*×*10^3^ images. As explained in the Results, the training and testing conditions were, respectively, the on-cross and off-cross stimuli shown in Figure 3a (red frames). In the experiments where the images from the study of Alemi-Neissi and colleagues^5^ were used, we trained all the SVMs on a mixture of unoccluded and partially occluded images, with the same probability of occurrence (50%) for each image type. While in ref.^5^ only a few selected object views were presented in their occluded version (marked by the red stars in Fig. 4b), the SVMs, during training, were exposed to partially occluded views of all the objects transformations included in the stimulus set (i.e., all variations in size, azimuth rotation, horizontal shift and in-plane rotation shown in the axes of Fig. 4b). As explained in the Results, this was necessary, because, otherwise, the generalization of the SVMs to the occluded views was very poor and did not allow recovering the saliency maps for all combinations of stimulus conditions and layers (see next section). For the accuracy analysis shown in Figure 4b, we tested separately 2.5 *×* 10^3^unoccluded and 2.5 *×* 10^3^ randomly occluded images. For inferring the saliency maps underlying the SVMs’ visual strategies, the number of test conditions is reported in the next section. Finally, in the experiments where the stimulus set of Djurdevic and colleages^12^ was used, we tested the SVMs with 2.5 *×* 10^3^ images separately for both the *regular* and *outline* object conditions (see Figure 7).

In the experiments with the stimulus set of ref.^5^, we additionally trained a pixel-based classifier in the object discrimination task originally administered to the rats (cyan curves/stars in Fig. 4b). Also in this case, we used a linear SVM classifier (using sklearn.svm) and trained it to predict the object labels using the full set of pixels (224 *×* 224) as input features. For consistency with the experiments done with VGG-16, we trained the pixel-based classifier with 5 *×* 10^3^ images, using an even mixture of unoccluded and occluded images sampled from the entire stimulus set, and we tested it with 2.5 *×* 10^3^ images separately for each image type.

All classification experiments were repeated 5 times (referred to, in the Results, as *classification runs*). In each run, we performed a full resampling of the subpopulations of units in each VGG-16 layer, whose activations were fed to the linear SVMs. In addition, in each run, all training and test images underwent different random augmentations.

### Visual Strategy Identification

We used two different approaches to infer the visual strategies used by the SVM classifiers, trained with VGG-16, layer-based activations. The first was based on the classification of partially occluded views of the visual objects; the second on the classification of random structural variations of a target object. Both approaches closely matched the image classification methods used, respectively, by Alemi-Neissi and colleagues^5^ and Djurdevic and colleagues^12^ to uncover rat visual perceptual strategies.

In the first approach, known as the *Bubbles method* ^55^, partially occluded images of each object view are obtained by imposing a random mask *m_ij_ ∈* [0, 1] on top of the image, with *i, j* being spatial indices indicating the pixel coordinate in the image. Each original pixel *p_ij_* is alpha-masked via 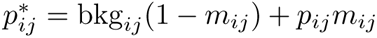, where we take bkg*_ij_* to be a uniform black background. The occluding mask *m_ij_* is constructed by randomly sprinkling *N_b_* Gaussians of fixed variance *σ* (the *bubbles*) in a region centered on the object and spanning 70% of the total image size. In our experiments, we fixed *N_b_* = 40 and set *σ* such that the resulting bubbles had a size equivalent to 2*^◦^* of visual angle. These values were selected based on the experimental design of Alemi-Neissi and colleagues^5^, who used bubbles of 2*^◦^* of visual angle, while *N_b_*was rat-specific and was chosen so as to bring each rat performance to be 10% lower than with the unmasked objects. Since *N_b_* ranged between 10 and 90, depending on the proficiency of the rat, in our tests with VGG-16 we set *N_b_* = 40, as an intermediate rat value. In our experiments, image masking occured before the random image augmentation described previously.

To derive the visual strategy of the observer for a particular object view, one has to compute the classification labels *ℓ_µ_∈* {0, 1} (where 0 represents an error and 1 represents a correct choice) for a set of masked images 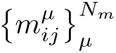, where *µ* is the index of the specific mask. Following ref.^5^, we took *N_m_* = 3000. The saliency map *S_ij_* obtained for the object view can then be reconstructed as:

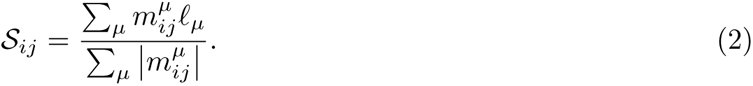

To assess whether the saliency value *S_ij_* computed for a given pixel was significantly correlated with the choices of the observer, we performed the following permutation test. We sampled 1000 random permutations of the classification vector, obtaining 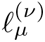, with *ν ∈* {0, 1*,…,* 1000}. We then applied eq. (2) using 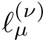 to estimate the null distribution 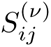. We then used a one-tailed statistical test to find which pixel values *S_ij_* were statistically higher than expected by chance (i.e., with *p < p*_th_) according to these null distributions 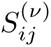, in such a way to identify significantly salient regions. Similarly, significantly anti-salient regions were identified as the sets of pixels for which *S_ij_*was statistically lower (i.e., with *p < p*_th_) than expected by chance. For a visual comparison among the saliency maps obtained for the rats and the SVMs trained on VGG-16 layer-based representations, we set *p*_th_ = 0.05, i.e., the same value used by Alemi-Neissi and colleagues^5^ (this threshold yielded the salient and anti-salient regions highlighted by the red and cyan patches in Fig. 5). Instead, for the overlap analysis shown in Fig. 6, *p*_th_ in the saliency maps obtained for the SVMs was independently adjusted, for every tested object view and each layer of the network, in such a way that the overall area of the salient region matched the one of the average saliency map obtained for the rat. This ensured that any difference in the amount of overlap between the maps obtained for different object views was not merely due to the different size of the salient regions being measured for the rats and the SVMs.

The second approach, introduced by Djurdevic and colleagues^12^, follows a different experimental paradigm: the subject is trained to recognize a given target object against a set of “distractor” objects. Once the classification is learnt, the subject enters a testing phase where is exposed to random shape-variations 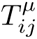 of the target object and its spontaneous choices *r_µ_ ∈* {0, 1} are collected (here 0 indicates that the random variant *µ* is classified as a distractor, while 1 indicates that is classified as being the target object). The saliency map associated to the target object can then be estimated (as illustrated in Fig. 7a) as:

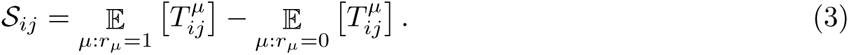

Also in this case, the statistical significance of the pixels in the saliency map was estimated via a permutation test. We produced 100 random permutations of the observer’s classification vector 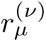, with *ν ∈* {0, 1*,…,* 100}, from which we estimated the corresponding null distributions of saliency values 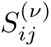, by applying eq. 3 using 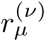. For each pixel, we then fitted a one-dimensional Gaussian to obtain a smooth version of the null-distribution. Finally, using the estimated Gaussian distribution, we performed a one-tailed test by comparing where the pixel values of the original saliency map (i.e., those obtained via the actual subject response vector *r_µ_*) sit with respect to the null distributions: a pixel was deemed significantly salient if it fell on the positive tail (*p <* 0.01), and significantly anti-salient if it fell on the negative one (*p <* 0.01).

## Acknowledgement

This work was supported by: the Italian Ministry of University and Research (D.Z.) under the calls PRIN 2022 (project no. 2022WX3FM5) and PRO3 (project NEMESI); the Simons Foundation Collaboration on the Global Brain grants 542989SPI and NC-GB-CULM-00002734 (A.A.); NIH Grant R01 NS104926 (A.A.). We thank Alessio Ansuini for easing access to data of ref.^12^ and Laura Porta for her contribution to initial, pilot tests with VGG-16. We also thank Gabriel Kreiman, Fabio Anselmi and Alessandro Sanzeni for valuable discussions and insightful feedback on our manuscript.

